# SarcAsM: AI-based multiscale analysis of sarcomere organization and contractility in cardiomyocytes

**DOI:** 10.1101/2025.04.29.650605

**Authors:** Daniel Härtter, Lara Hauke, Til Driehorst, Yuxi Long, Guobin Bao, Andreas Primeßnig, Branimir Berečić, Lukas Cyganek, Malte Tiburcy, Christoph F. Schmidt, Wolfram-Hubertus Zimmermann

## Abstract

Cardiomyocyte function critically depends on sarcomere dynamics and their organization in myofibrils. To uncover how cardiomyocyte function emerges from individual sarcomere dynamics, comprehensive analysis of Z-bands (nanometer scale), sarcomeres (∼2 µm scale), and myofibrils (∼10 to 100 µm scale) is required. Tools for such multiscale analyses are presently lacking. Here we introduce the Sarcomere Analysis Multitool (SarcAsM), which combines deep learning and graph-based methods for automated, fast, and unbiased structural assessment of sarcomeric Z-bands, sarcomeres, and myofibrils as well as their organization in larger myofibril domains. SarcAsM features a generalist deep learning model, pre-trained on a broad range of experimental and published images, ensuring its versatile application and immediate usability across diverse datasets. Finally, we demonstrate the versatile utility of SarcAsM in analyzing sarcomere structure and dynamics under acute and chronic drug exposure in hiPSC-derived cardiomyocytes with fluorescently tagged Z-bands. SarcAsM is available as open-source Python package and stand-alone application.

---

Cardiac muscle contraction emerges from the collective dynamics of billions of cardiomyocytes^1^ each containing ∼100 million myosin motor heads^2–4^ organized into thick filaments. At the subcellular level, these thick filaments interdigitate with actin filaments and form sarcomeres, the ∼2 µm long basic uniaxial contractile muscle units. In cardiomyocytes (∼100 µm long), up to 50 sarcomeres can be connected in series to form a myofibril. The parallel alignment and network connectivity of myofibrils are essential for coordinated contractile force transmission in cardiomyocytes. In healthy adult cardiomyocytes, sarcomeres and myofibrils are uniformly registered, whereas sarcomere and myofibril disarray are hallmarks of heart failure^5,6^.

The advent of human pluripotent stem cells (PSCs)^7,8^ coupled with robust protocols for directed cardiomyocyte differentiation and culture^9,10^, and methods for stably inserting fluorescent reporters into sarcomere proteins^11–13^ are opening the door to detailed longitudinal studies of sarcomere structure and function – providing new opportunities for drug development and disease modeling^14–16^. Plated on micropatterned substrates, PSC-derived cardiomyocytes assume *in-vivo*-like geometries^17–21^ with anisotropic myofibril assemblies, making it possible to further standardize sarcomere analyses. Combined with automated microscopy techniques, this set of novel experimental tools enables high-throughput experiments, creating the challenge to process the resulting massive data in a fast, robust, and unbiased manner.

Several computational tools have been introduced to analyze aspects of cardiomyocyte and sarcomere structure and function^22–29^. Some tools focus on Z-band morphology (ZlineDetection^23^ or sarcApp^29^). Other tools investigate sarcomere function by quantification of sarcomere lengths and motion using Fourier transform (SarcOptiM^24^), by measuring the distance between the centroids of adjacent Z-bands (Sarc-Graph^26,30^), or by wavelet analysis (SarcTrack^22^). The existing tools, with the exception of sarcApp^29^, build on classical image processing approaches for detecting Z-bands or sarcomeres that require high signal-to-noise data with clearly distinguishable Z-bands. This can lead to selection bias of well-structured over disordered morphologies, which may severely limit applications, in particular in drug screens. Robust AI-based deep learning tools enabling multiscale structural and functional analysis of sarcomeres—from Z-bands to myofibrillar organization—and rapidly and robustly processing diverse data sets (images/movies) across experimental models (patterned/unconfined cultures, stained cells/tissues, reporter models) have been lacking.

To close this gap, we designed SarcAsM (Sarcomere Analysis Multitool) as a fast, robust, accessible, and user-friendly computational tool suitable for a wide variety of studies of sarcomere structure and function, including morphogenesis, disease phenotyping, biophysical studies of contractility, and drug testing. SarcAsM employs a neural network to simultaneously segment masks for sarcomere Z-bands, sarcomere regions, and cells, and to estimate M-band masks and sarcomere orientations. Based on this information, SarcAsM determines a comprehensive set of structural features at the Z-band, sarcomere, myofibril, and cell level in an efficient, unbiased, and quantitative manner. For high-speed recordings of beating cardiomyocytes, SarcAsM automatically identifies lines of interest (LOIs) along myofibrils to track and analyze the motion of individual Z-bands and sarcomere lengths with nanometer scale spatial resolution. In conjunction with a custom-generated sarcomere reporter (ACTN2-Citrine) induced pluripotent stem cell (hiPSC) model for stable sarcomere labeling in cardiomyocyte derivatives, we demonstrate the capabilities of SarcAsM to precisely quantify (1) structural effects of chronic drug exposure of cardiomyocytes in 2D monolayer culture and (2) functional effects of acute drug administration on individual cardiomyocytes cultured on micropatterned soft gels. SarcAsM is available as a stand-alone application with a graphical user interface (GUI) and as a Python package with a high-level application programming interface (API) for customization and integration into analysis pipelines (https://sarcasm.readthedocs.io/en/, https://github.com/danihae/SarcAsM).

## Results

### Endogenous fluorescent labeling of sarcomeres using CRISPR/Cas9

For live-cell imaging of hiPSC-derived cardiomyocytes, we developed a sarcomeric Z-band reporter hiPSC-line by fusion of the C-terminus of the endogenous alpha-actinin-2 (*ACTN2*) gene with the fluorescent protein Citrine^31,32^ using CRISPR/Cas9 (**Fig. S1a** and **Table S1**). We differentiated hiPSCs into cardiomyocytes using an established protocol and confirmed unimpaired contractility of the ACTN2-cardiomyocytes in a well-established engineered heart muscle (EHM) assay^10^ (**Fig. S1b-e**).

### Detection of sarcomere Z-bands, M-bands, orientation field and cell/sarcomere mask using deep learning

The alpha-actinin-2 protein, with its N-terminal actin-binding domains, is localized not only in sarcomere Z-bands but also in dense bodies during sarcomerogenesis and at cell adhesion sites^33^, creating ambiguity in Z-band detection. Additionally, the fluorescence signal of Z-bands is naturally non-uniform (**Fig. 1a**) due to structural variability and imaging noise. To address these challenges, we employ a deep learning approach using a U-Net++ convolutional neural network^34^, which robustly detects sarcomeres as distinct parallel band structures while effectively excluding “off-target” signals such as stress fibers (**Fig. 1 a,b**). Our U-Net++ model simultaneously predicts multiple features from the input images: cell and sarcomere masks (**Fig. 1b**), sarcomere orientation field (**Fig. 1c**), and Z-band and M-band segmentations (**Fig. 1d**). Notably, the sarcomere orientation field is a 2D vector field pointing from M-bands towards Z-bands, providing crucial information about sarcomere directionality (**Fig. 1c**).

**Figure 1:**
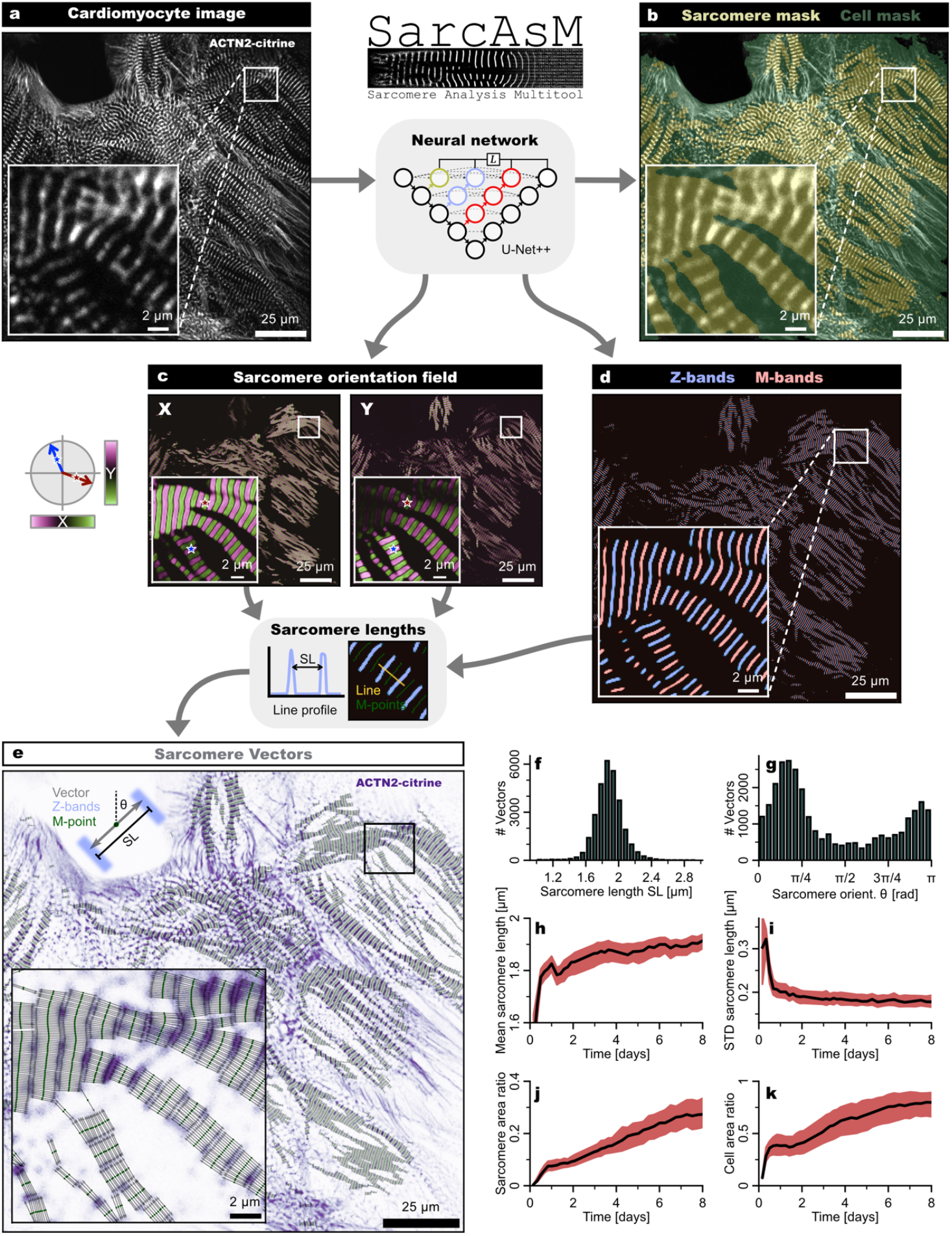
Deep learning-based sarcomere segmentation and spatial mapping of sarcomere length and orientation using sarcomere vectors. (**a**) Live confocal microscopy image (z-slice with max. intensity) showing hiPSC-derived cardiomyocytes (culture day 6) with endogenous ACTN2-citrine fluorescence labeling of sarcomeric Z-bands. Inset highlights the sarcomere structure at higher magnification. A multi-headed U-Net++ deep learning model simultaneously predicts multiple image features trained on expert-annotated data: (**b**) Binary masks of cells (green) and sarcomeres (yellow). (**c**) Sarcomere orientation field composed of unit vectors representing the direction from M-band to Z-band within each sarcomere. At a given pixel, the field maps the local orientation of sarcomeres. The X- and Y-components of unit vectors are shown as color-coded maps with insets providing higher magnification views to highlight the detailed spatial precision of the orientation field. (**d**) Z-bands (blue) and M-bands (red). The segmentation accurately identifies Z-bands and further estimates the position of M-bands despite signal non-uniformity, with minimal false positives or misclassifications and exclusion of non-sarcomeric features such as stress fibers. (**e**) Sarcomere vectors (grey arrows) representing local sarcomere length and orientation. M-points (green) are extracted along M-bands, and sarcomere orientation is derived from the orientation field. Sarcomere length is measured as the distance between Z-band peaks in a profile line along the orientation vector. The inset shows a magnified view, illustrating detailed and accurate mapping of local sarcomere structure. (**f, g**) Histograms showing the distributions of sarcomere lengths (SL) and orientations (θ) within the analyzed image shown in panel **a**. (**h**) Time-course of mean sarcomere lengths of cardiomyocytes in unconfined 2D culture. (**i**) Time-course of standard deviation of sarcomere lengths. For both (**h**) and (**i**): Black line and red interval show median and interquartile ranges of 216 fields of views (FOVs) in 24 wells, weighted by sarcomere area ratio (single FOV shown in panels **a-d**). (**j**) Time course of sarcomere area ratio (percentage of cell area containing sarcomeres) of cardiomyocytes in unconfined 2D culture. (**k**) Time course of cell area ratio (percentage of FOV area occupied by cells). For both (**j**) and (**k**): Black line and red interval show mean and STD of 216 FOVs (215×215 µm) in 24 wells.

This multi-target prediction approach allows for comprehensive analysis of sarcomere structure and organization. The training data for our model was compiled from diverse sources, including our own data and publicly available datasets, spanning different cell types, labeling strategies (e.g., endogenous labeling and immunostaining), and microscopy modalities. This diverse dataset, enriched through extensive data augmentation, enables the training of a generalist model applicable to a wide range of experimental data (**Table S2, Fig. S2**). The ground truth for training was generated through a multi-step approach combining computationally intensive custom double wavelet analysis^22^ and manual annotation and curation (details provided in Methods and **Supplementary Note 1**). We benchmarked the detection of sarcomeres against manually annotated ground truth in a diverse dataset, achieving excellent accuracy and specificity with an F1 score of 0.86 (details see **Supplementary Note 2**).

For robust Z-band localization in high-speed movies of beating cardiomyocytes, we employ a time-consistent convolutional neural network architecture (3D-U-Net)^35,36^ that predicts Z-bands not just frame-by-frame, but incorporates information from preceding and subsequent frames to improve frame-to-frame consistency and reduce flickering artifacts due to imaging noise (**Movie 1, Fig. S3a-d**), enhancing the temporal stability of Z-band predictions in dynamic cardiomyocyte recordings.

### Spatial mapping of sarcomere lengths and orientations using sarcomere vectors

To precisely assess local sarcomere length and orientation, we developed a novel approach using “sarcomere vectors” based on the outputs of our deep learning model. We first extracted a dense set of points along the M-bands using thresholding and skeletonization techniques. At each of these points, the local sarcomere orientation was determined from the orientation field (**Fig. 1c**). To measure local sarcomere lengths, we analyzed the Z-band intensity profile along a line with sarcomere orientation at each point. The distances between Z-band peaks were calculated with sub-pixel accuracy. This procedure results in a large set of ‘sarcomere vectors’, akin to compass needles, accurately reflecting local sarcomere location, length and orientation (**Fig. 1e**). This dense ‘field’ of sarcomere vectors is a highly detailed and truthful local representation of sarcomeres, allowing us to quantify the distribution of sarcomere lengths and orientations in a given image (**Fig. 1f,g**). To validate our approach, we benchmarked the sarcomere length analysis against manually measured sarcomere lengths in a diverse set of images. SarcAsM demonstrated excellent accuracy in both the average length (mean deviation of 19 nm) and standard deviation per image (mean deviation of 22 nm) across all test images (details see **Supplementary Note 2**).

We cultured ACTN2-citrine cardiomyocytes in a 96-well plate format for 8 days with fluorescence image acquisition from defined positions every 4 hours using a Yokogawa CV8000 high-throughput automated imaging station. In each image we then analyzed the sarcomere vectors. The sarcomere length increased slightly within physiological values (**Fig. 1h**), while the variation of sarcomere lengths, measured by the standard deviation in each image, decreased strongly between day 0-2, indicating sarcomere maturation (**Fig. 1i**). Both the area occupied by sarcomeres (**Fig. 1j**), normalized by cell area, and the area occupied by cells in each image (**Fig. 1k**) increased steadily over 8 days of cardiomyocyte monolayer culture, which is in line with hypertrophic growth.

### Analysis of Z-band morphology, assembly, and abundance

To analyze the morphology of Z-bands, we segmented individual Z-bands using thresholding and labeling of the Z-bands predicted by deep learning (**Fig. 2a,b**). For each Z-band object, we quantified several features, including length (**Fig. 2c**), straightness (**Fig. 2d**), and the fluorescence intensity per length (**Fig. 2e**), as a measure of ACTN2 protein abundance. The analysis of the 8-day movies revealed a steady increase in Z-band lengths (**Fig. 2f**) along with a decrease in Z-band straightness (**Fig. 2g**), and an increased ACTN2 abundance at the Z-band level after culture day 4 (**Fig. 2h**). Taken together, these parameters precisely quantify the Z-band assembly and maturation in extended (8 day) cardiomyocyte cultures in a 96-well culture format.

**Figure 2:**
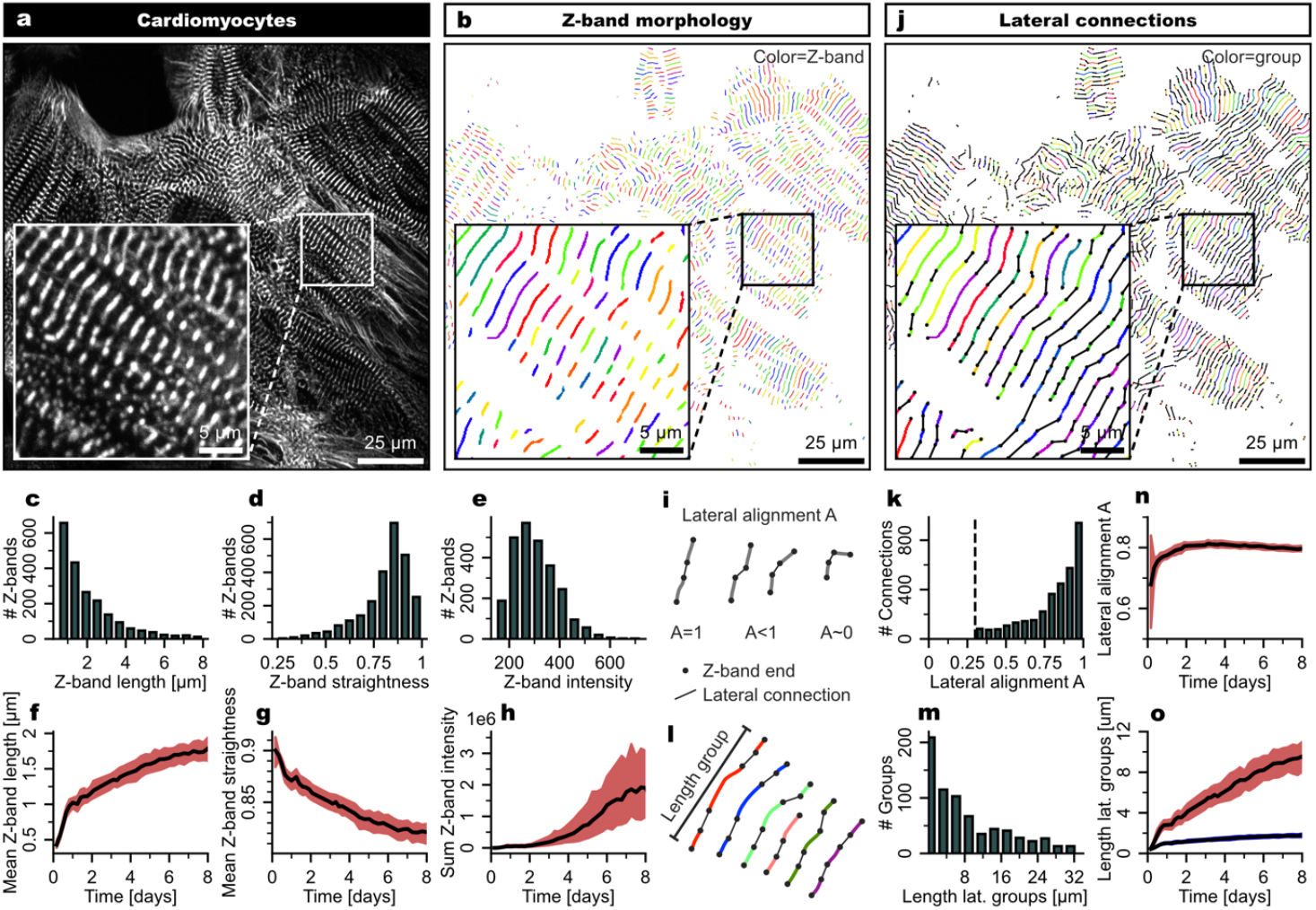
Analysis of individual Z-band morphology and lateral Z-band organization. (**a**) Live confocal microscopy image (z-slice with max. intensity) of hiPSC-derived cardiomyocytes (culture day 6) in unconfined 2D culture with endogenous fluorescent tagging of alpha-actinin-2 (ACTN2) to label the sarcomeric Z-bands. Inset shows magnification. (**b**) Object-based segmentation of Z-bands. Colors represent object labels. (**c-e**) Distributions of Z-band lengths, straightness (1=straight, <1=curved) and intensities in single image shown in **a**. (**f-h**) Time-course of Z-band length, straightness and total Z-band intensity of cardiomyocytes in unconfined 2D culture. (**i**) Schematic illustration of the lateral alignment metric AA, which quantifies the alignment of neighboring Z-bands. (**j**) Analysis of lateral organization of Z-bands using a linear sum assignment algorithm to identify lateral connections between neighboring Z-bands. Connections are only established for Z-band ends with alignment >0.3 and within 4 µm distance. Black dots mark Z-band ends, black lines indicate lateral connections, and matching colors represent groups of laterally connected Z-bands. (**k,m**) Distributions of lateral alignment A of lateral connections, and total length of groups of laterally connected Z-bands. (**n,o**) Time-course of lateral alignment A, and total length of groups of laterally aligned Z-bands (vs. individual Z-band length depicted in blue) of cardiomyocytes in unconfined 2D culture. The xy-plots show medians and quartiles of feature averages in 216 fields of view (215×215 µm) in 24 wells, weighted by sarcomere area ratio.

### Analysis of higher-order spatial organization of sarcomere Z-bands

To gain insight into the higher-order Z-band organization, we identified the location and orientation of the two ends of each Z-band and calculated pairwise Euclidean distances *D* between all Z-band ends. For Z-band pairs within a 4 µm distance, we also computed a custom lateral alignment metric *A*, which quantifies the degree of alignment between neighboring Z-bands (**Fig. 2i, Fig. S4**). Using these metrics, we applied a linear sum assignment algorithm to optimally connect Z-band ends, ensuring that the total lateral misalignment (1*−*A) was minimized across all pairs (**Fig. 2j**). This approach allowed us to group laterally connected Z-bands into networks, providing insights into their spatial organization and alignment. Based on this data, we could quantify the distributions of lateral alignment *A* (**Fig. 2k**), and the total length of groups of laterally aligned Z-bands (**Fig. 2l,m**). In long-term cardiomyocyte culture, increasing alignment (**Fig. 2n**) and growth of lateral groups that is more than proportional to the lengths of individual Z-bands (**Fig. 2o**), suggests highly ordered concentric hypertrophic growth.

### Automated identification of myofibrils and cell-level myofibril domains

To identify and quantify cardiomyocyte sarcomere assemblies in unconfined isotropic monolayer cultures, we estimated myofibril positions, lengths, and curvatures based on the set of sarcomere vectors. Starting from a randomly selected subset of, e.g., 10% of sarcomere vectors, we grew lines into both directions connecting series of points using a nearest neighbor algorithm (**Fig. 3a**). Line growth stopped when there were no further points in proximity. Short lines with less than 5 sarcomere vectors were discarded. The resulting lines precisely sampled myofibrils (**Fig. 3a**) and, after transformation into a spatial myofibril length map (**Fig. 3b**), allowed us to quantify the length distribution of myofibrils in each image (**Fig. 3c**). We also calculated the bending (mean squared curvature) of each line, with large values reflecting highly curved or wiggly myofibrils (**Fig. 3d**). The observed increase in myofibril lengths with decreasing bending energy, i.e., increasing straightness, over time in culture (**Fig. 3e,f**) was again consistent with hypertrophic cardiomyocyte growth and structural maturation.

**Figure 3:**
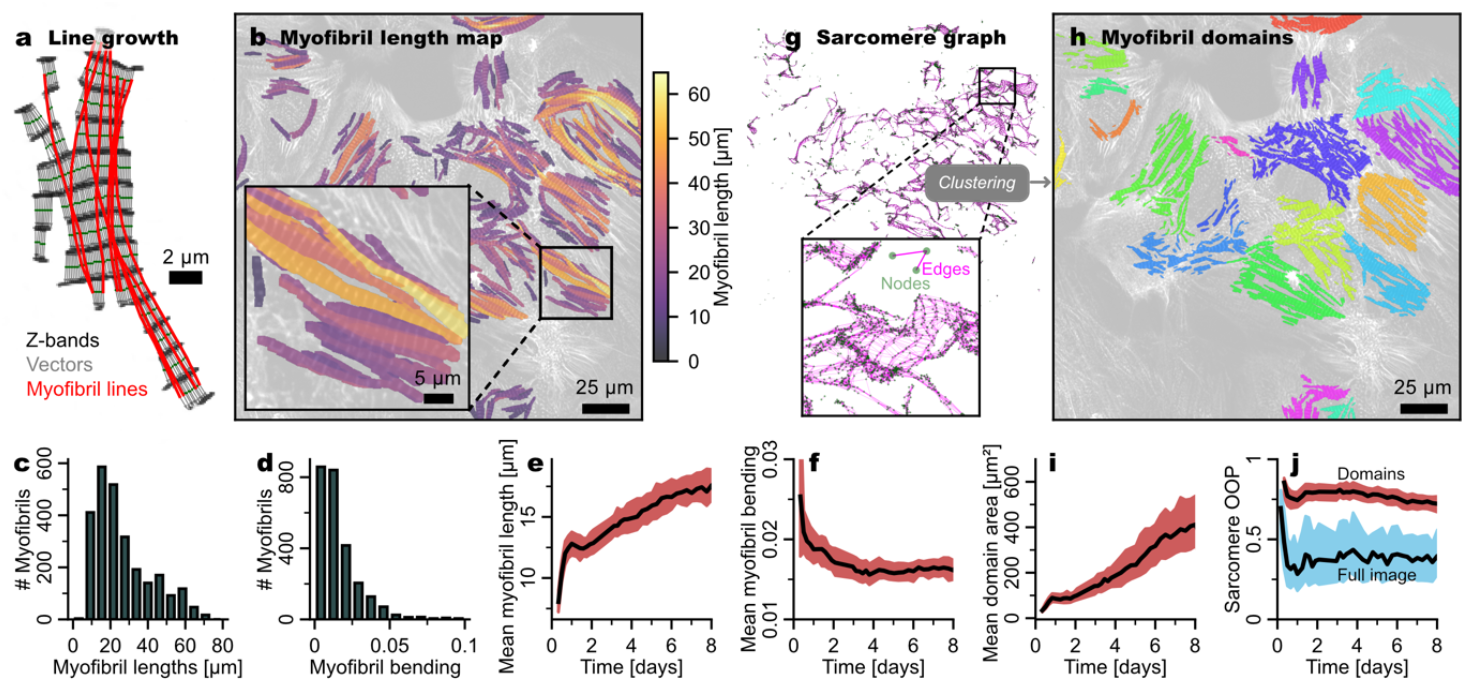
Myofibril line growth algorithm and clustering of myofibril domains. (**a**) Illustration of sarcomere line growth algorithm with myofibril lines (red) starting from randomly selected seeds (e.g., 10% of sarcomere vectors). Green points show midline points and grey arrows sarcomere vectors. Z-bands are shown in black. (**b**) Spatial map of myofibrils lengths computed from set of myofibril lines. Color represents the mean length of myofibrils. (**c**) Histogram of myofibril line lengths from the data in **b**. (**d**) Histogram of bending of myofibril lines of data in **b**, quantified by the mean squared curvature of lines. (**e,f**) Time course of myofibril lengths and bending of cardiomyocytes in unconfined 2D culture. (**g**) Sarcomere graph constructed from a set of sarcomere vectors. Edges were added between sarcomere vectors in proximity and with angular similarity. The network representation was created using the Fruchterman-Reingold force-directed algorithm with 50 iterations. (**h**) Myofibril domains identified by the graph-based Leiden community detection algorithm. Sarcomeres of one domain are marked by the same color. (**i**) Time-course of myofibril domain areas in unconfined 2D culture. (**j**) Time-course of sarcomere orientation order parameter *OOP* within myofibril domains (red), representing for the most part individual cardiomyocytes. The average *OOP* for the full field of views is included for comparison (blue), demonstrating that individual cardiomyocytes (indicated by myofibril domains) were not aligned. **e,f,i,j** show median and interquartile range of 216 field of views (215×215 µm) in 24 wells of extended (8 day) unconfined 2D cardiomyocyte culture.

The set of sarcomere vectors reflecting local sarcomere length and orientation was further used to identify spatial domains of interconnected and similarly oriented sarcomeres, reflecting higher order contractile units essential for coordinated and forceful contractions at the cell-level. We created a graph from the set of sarcomere vectors, with edges between sarcomere vectors of the same and adjacent sarcomere midlines, and used the graph-based Leiden community detection algorithm^37^, a heuristic method based on modularity optimization, to partition the sarcomere graph into domains (**Fig. 3g,h**). During long-term cardiomyocyte monolayer culture, we observed a steady increase in highly ordered domains (**Fig. 3i**), indicating coordinated cardiomyocyte growth with improved cell-scale sarcomere order. Of note, the orientational order of sarcomeres within domains, quantified by the orientational order parameter (OOP), was markedly higher than that of full images (**Fig. 3j**).

### Long-term morphological phenotyping under chronic drug treatment

To test whether SarcAsM can be applied to identify sarcomere targeted drug activities, we applied the cardiac myosin inhibitor Mavacamten from day 3 of cardiomyocyte culture for 5 consecutive days at clinically relevant concentrations of 0.1 - 10 µM (**Fig. 4a-d, Movie 2**). Mavacamten elicited strong concentration-dependent effects: cardiomyocytes growth increased already at low concentrations (**Fig. 4e**), whereas the cardiomyocyte area occupied by sarcomeres decreased at higher concentrations (**Fig. 4f**); sarcomere structures were substantially altered within hours after Mavacamten administration in a concentration-dependent manner (**Fig. 4g-m**). In particular, Z-band lengths, the total length of laterally aligned groups of Z-bands and lateral alignment decreased (**Fig. 4g-i**). Average sarcomere length increased slightly at low concentration, which may be the result of reduced passive tension generated by myosin - consistent with the mode of action of Mavacamten - while strongly decreasing at high concentration to levels comparable to those observed after cell plating at day 0 (**Fig. 4j**). At medium and high concentrations, the variance of sarcomere lengths showed a strong transient (1 µM Mavacamten) and sustained (10 µM Mavacamten) increase (**Fig. 4k**), while myofibril lengths and myofibril domain sizes decreased strongly (**Fig. 4l,m**). We calculated the multivariate Z-factor to above 0.8 for higher concentrations 1 day after drug addition, indicating excellent assay quality, confirming SarcAsM’s suitability for high-throughput screening with single-replicate measurements. These observations are particularly interesting in light of the clinical use of doses resulting in plasma levels of 350-700 ng/mL (1.25-2.5 µM) and clinically reported unwanted side effects such us heart failure exacerbation in patients with hypertrophic cardiomyopathy under Mavacamten treatment^38^.

**Figure 4:**
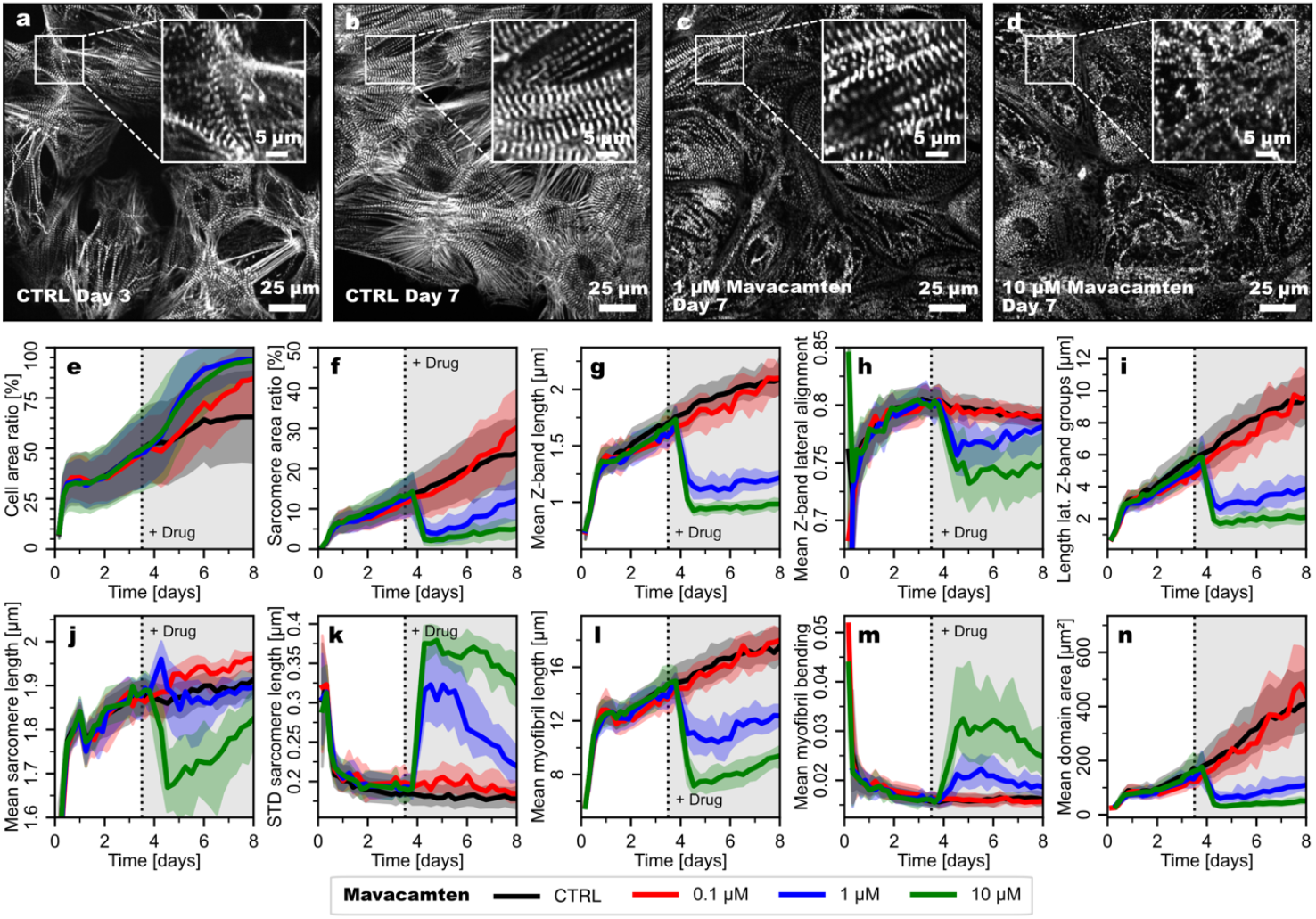
Effect of exposure to the myosin-inhibitor Mavacamten at clinically relevant concentrations. Representative images (**a**) on culture day 3 before treatment, (**b**) on culture day 7 without treatment (control), (**c**) on culture day 7 in the 1 µM Mavacamten treatment group (4 days of Mavacamten), and (**d**) on culture day 7 in the 10 µM Mavacamten treatment group (4 days of Mavacamten). (**e**-**n**) Time course of sarcomere structural features in control and Mavacamten-treated cardiomyocytes. For each condition, 8 wells with 9 FoVs (215×215 µm) each were recorded (single FoVs shown in **a**-**d**). Curves show median and interquartile ranges of FoVs, weighted by sarcomere area ratio per FoV for **g-n**. Gray area indicates time interval of drug treatment.

### Individual cardiomyocytes on micro-patterned soft gels for sarcomere motion analysis

To assess the contractile function of isolated beating cardiomyocytes, we cultured ACTN2-citrine CMs on micropatterned polyacrylamide hydrogels with viscoelastic properties and shapes that mimic healthy myocardium^39^ (micropattern patch size 12×70 µm, Young’s modulus 15 kPa). In contrast to unstructured 2D culture, cells in this assay format adapt to the shape of the micropattern, developing ordered myofibrils and greatly reducing cell-cell variability in shape and sarcomere structure^17,40^. Furthermore, the assay allowed us to quantify and compare the contractility of cells due to the well-defined elastic environment. After 20 days of maturation, the motion of isolated spontaneously beating cardiomyocytes was recorded with a confocal microscope at a high frame rate (66 frames per second) for 20 seconds (**Fig. 5a, Movie 1**). In this way, we could rapidly measure the motion of ∼1,800 individual cells and ∼10,000s sarcomeres, yielding sufficient statistics to sample the biological variability and resolve even subtle effects.

**Figure 5:**
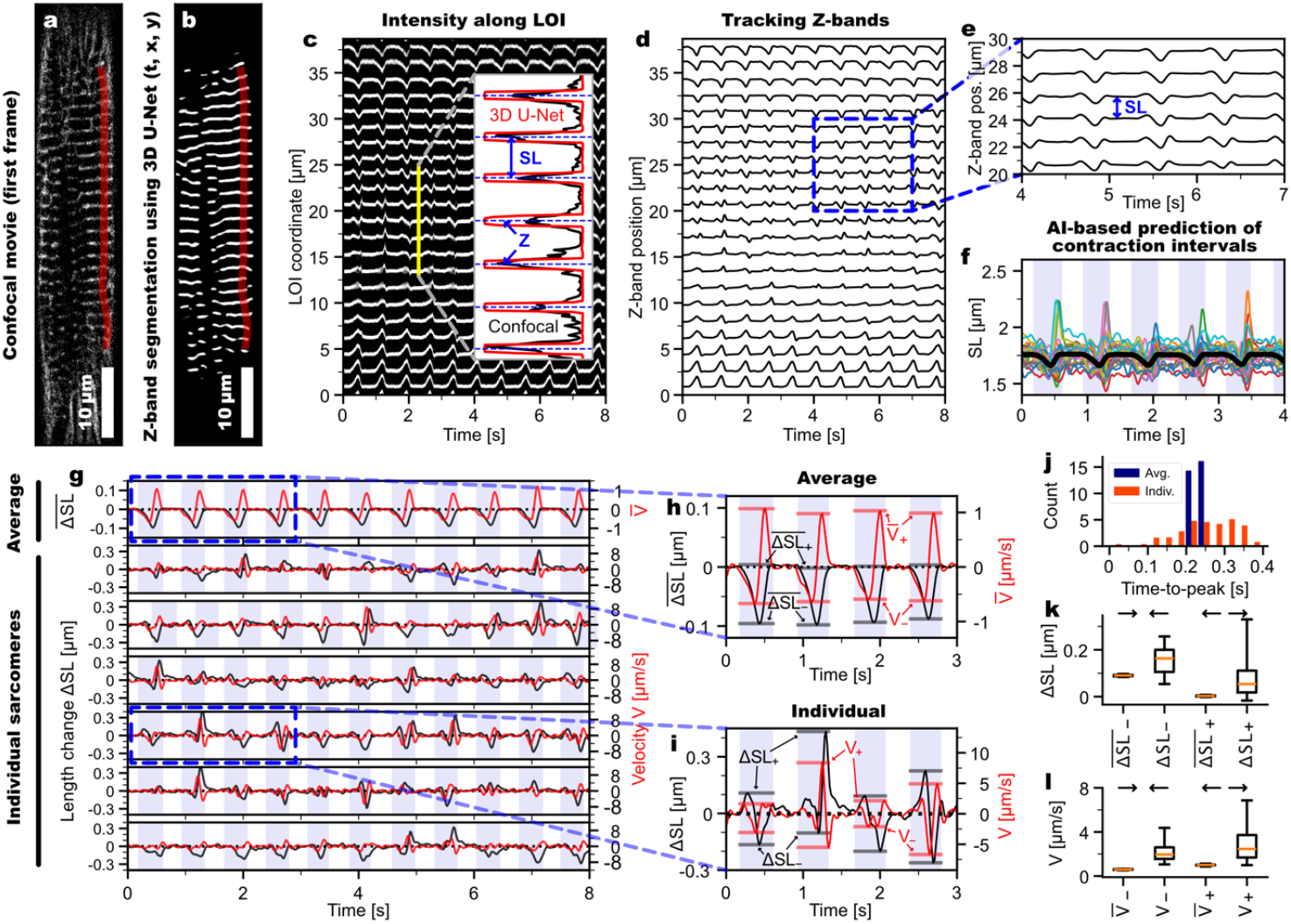
Tracking and analysis of individual and average sarcomere motion in cardiomyocytes on a micropatterned soft gel. (**a**) Confocal image from a 1,400-frame recording of a representative spontaneously beating cardiomyocyte (66 frames per second). (**b**) Sarcomere Z-bands detected with time-consistent convolutional neural network (3D-U-Net). LOI is displayed in yellow. (**c**) Kymograph of Z-bands motion along automatically detected LOI in panel **b** with line width 10 pixels/0.72 µm. Inset shows the intensity profile along the LOI at the single time point displayed in panel **c**. Black line shows raw microscopy data, red line the deep learning result. (**d**-**e**) Individual Z-band trajectories *Z*(*t*). (**f**) Time series of sarcomere lengths *SL*(*t*) (colored=individual, black=average). The blue shaded background shows contraction time intervals predicted by a custom convolutional neural network. (**g**) Top row: average sarcomere length change 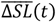 (black) and velocity 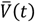 (red); bottom row: individual sarcomere length changes Δ*SL*(*t*) (black) and velocities *V*(*t*) (red), selection of sarcomeres from **d**. (**h**-**i**) Enlargement of average and individual sarcomere motion in panel **g** with illustration of quantification of minimum and maximum length changes and velocities within each contraction cycle. (**j**) Histogram of time to peak *T*_*p*_ for individual and average sarcomere length change. (**k**) Boxplot of minimum (*−*) and maximum (+) sarcomere length changes Δ*SL*(*t*). (**l**) Boxplot of minimum (−) and maximum (+) sarcomere velocities *V*(*t*). Boxes show quartiles, red lines the median, and whiskers the 5th and 95th percentiles of the distributions. **j**-**l** show data from one representative LOI in the cardiomyocyte in **a.**

### Automated identification of regions suitable for sarcomere motion tracking

To quantify sarcomere motion in a fast, automated, and unbiased manner, we created a custom algorithm to automatically identify lines along well-ordered myofibrils suitable for tracking sarcomere motion. We adapted the myofibril line growth algorithm described above and created a set of 200 lines each with more than 10 sarcomeres in length and tuned for higher straightness (persistence=5; **Fig. S5a**). Because this set often contained multiple nearly identical lines, we identified groups of distinct lines using agglomerative clustering based on the pair-wise Hausdorff distances between lines (**Fig. S5b,c**). Finally, we selected the longest line of each cluster and selected the two longest lines as lines of interest (LOI) in each cell for sarcomere motion tracking (**Fig. S5d**). Alternatively, SarcAsM allows to randomly select LOIs from the initial set of lines. This automated line identification method is robust and applicable to diverse cardiomyocyte image datasets, including standard 2D cultures (**Fig. S5e-g**)

### Tracking and analysis of average and individual sarcomere motion

Assessing sarcomere motion based on the set of sarcomere vectors is limited by the length steps of the wavelet bank (e.g., 50 nm), making high precision computationally expensive. To analyze sarcomere motions, we have therefore extracted intensity kymographs along automatically identified LOIs (line width 0.72 µm) in the movies of contracting cardiomyocytes with Z-bands predicted by the time-consistent convolutional neural network (**Fig. 5b,c, Movie 1**). This procedure provided more precise and noise-insensitive localization of Z-bands than using raw confocal data. We then determined the exact location of Z-bands from the intensity profiles of the kymograph slices at each time point (**Fig. 5c inset**) and ultimately linked Z-band center positions across images into time series using an established tracking algorithm^41^ (**Fig. 5d,e, Movie 3**). The spatial resolution of our Z-band tracking method was ∼18 nm (**Fig. S3g**), confirmed by simulated data (**Movie 4**). To further smooth the time series *Z*(*t*), we applied a Savitzky-Golay filter (window length 11 frames, polynomial order 5), which preserves extreme amplitudes and does not introduce oscillatory artifacts. We calculated individual sarcomere lengths *SL*(*t*) by subtracting the positions of neighboring Z-bands, and from that the average sarcomere length 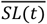 (**Fig. 5e,f**).

To relate individual sarcomere motions to the different phases of the cells’ contraction cycles, we predicted the exact start and end of each contraction and the intervening quiescent phase using a custom convolutional neural network (ContractionNet; **Fig. S6**) trained with expert-annotated and simulated training data (**Fig. 5f**). The inputs to the network are the time series *Z*(*t*) and *SL*(*t*) of all sarcomeres and the output is a binary time series *S*(*t*) with value 1 during contractions and value 0 during quiescent phases (**Fig. S6a**). “Contraction” denotes the whole duration of activity, including shortening and lengthening of individual sarcomeres during a contraction cycle. We found this method to be more sensitive and robust to noise than threshold-based methods, especially for small contraction amplitudes (**Fig. S6b**). Based on *S*(*t*), we calculated the spontaneous beating rate *BR*, the beating rate variability (STD of full cycle durations), and the durations of contractions *T*_*C*_ and quiescent phases *T*_*Q*_. Further, we determined the equilibrium length *SL*_0_of each sarcomere as the average length during the quiescent phases at *S* = 0. For each sarcomere, we calculated the sarcomere length change Δ*SL*(*t*) = *SL*(*t*) *− SL*_0_and, by temporal differentiation, the velocity *V*(*t*) (**Fig. 5g**). Finally, we calculated sarcomere average length change 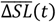 and average velocity 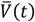(**Fig. 5g**). This analysis showed that the motion of individual sarcomeres was highly dynamic and heterogeneous and differed from contraction cycle to contraction cycle, whereas the average sarcomere motion was smooth and persistent over time.

Within each contraction cycle, we quantified the maximum and minimum length change Δ*SL*_−_ /Δ*SL*_+_ and the maximum and minimum velocity *V*_−_/*V*_+_ of individual sarcomeres and of the average (**Fig. 5h,i**). We also determined the time to peak *T*_*P*_, i.e., the time from the onset of the contraction to the maximal length change, for individual sarcomeres and the sarcomere average (**Fig. 5j**). For the average, the variance of *T*_*P*_ between contraction cycles was small, whereas for the individual sarcomeres we observed a broad distribution of *T*_*P*_, i.e., the individual sarcomeres were maximally shortened at different times during the contraction. Because of this asynchrony, the maximal activity of individual sarcomeres, both in length change and velocity, far exceeds the average activity, by a factor of ∼2 for ΔSL and by a factor of ∼5 for shortening (*V*_−_) and lengthening (*V*_+_) velocity (**Fig. 5k,l**). The average sarcomere length never extended beyond resting length, while individual sarcomeres frequently elongated beyond their resting length during all phases of contraction, but particularly at the end of contractions (a phenomenon of unknown pathophysiological relevance in cardiac muscle, termed sarcomere “popping”; **Fig. 5g,i**).

### Individual and average sarcomere dynamics under acute drug treatment

Similar to the chronic drug exposure experiment in unconfined monolayer culture, but now with a focus on drug-induced changes of contractility, we performed acute drug exposure experiments with reference compounds with well-characterized effects on cardiomyocyte contractility: Isoprenaline (β adrenoceptor agonist), Levosimendan (calcium sensitizer), Mavacamten (myosin inhibitor), Omecamtiv mecarbil (myosin activator), Verapamil (calcium channel blocker), and Digitoxin (sodium-potassium ATPase inhibitor). For each condition, we recorded spontaneously beating individual cardiomyocytes adherent to 15 kPa elastic substrates, followed by tracking and analysis of sarcomere motion in up to 4 automatically detected LOIs per cell (> 10 sarcomeres, > 10 contraction cycles; **Fig. 6a-d**). Arrhythmically beating cells (beating rate variability > 0.25 s) were excluded. We classified the drug effects into 3 categories: chronotropic, inotropic, and lusitropic (**Fig. 6e-j**, detailed data see **Fig. S7**). The observed effects at the average sarcomere level are consistent with the expected activities and modes of action of the reference compounds^42^. Interestingly, individual sarcomere motion patterns strongly differed from drug to drug and with drug concentration (**Fig. 6a’-d’**).

**Figure 6:**
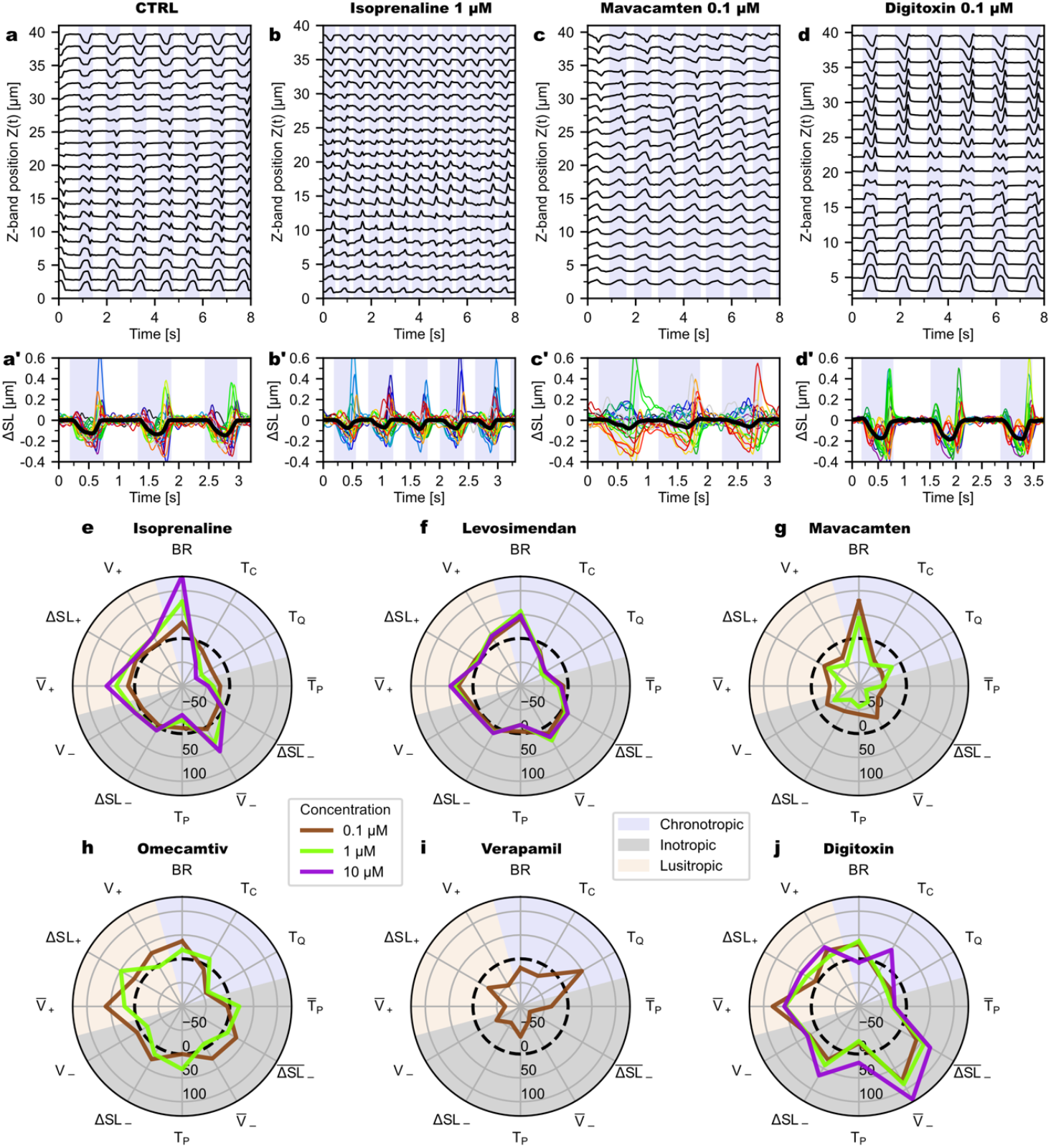
Effect of acute drug treatment on individual and average sarcomere motion. (**a**)-(**d**) Sarcomere Z-band trajectories of representative LOIs of an untreated cardiomyocyte (**a**) and drug-treated cardiomyocytes (**b**-**d**). (**a’**)-(**d’**) Overlay plots of sarcomere length changes of LOIs in (**a**)-(**d**). Thin colored lines show individual sarcomere motions, the black line shows the average motion. Blue-shaded regions mark contraction time periods (T_C_). (**e**)-(**j**) Radar charts of drug-treatment results, showing relative effect as change from median to median in % compared to untreated control (dashed circle). Missing conditions (10 µM Mavacamten, 1-10 µM Verapamil), because no spontaneously contracting cells were found. In total, 1,815 spontaneously beating cardiomyocytes were recorded and 6,944 LOIs were automatically detected and analyzed. For each condition, at least 20 LOIs with at least 10 contraction cycles and 10 sarcomeres were evaluated. Features: *BR*=beating rate, *T*_*C*_=duration contractions, *T*_*Q*_=duration quiescent phases, 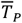=time to peak of sarcomere average, 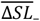=maximal shortening of sarcomere average, 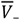=maximal shortening velocity of sarcomere average, *T*_*P*_=time to peak of individual sarcomeres, Δ*SL*_−_=maximal shortening of individual sarcomeres, *V*_−_=maximal shortening velocity of individual sarcomeres, 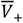=maximal elongation velocity of sarcomere average, Δ*SL*_+_=maximal elongation of individual sarcomeres, *V*_+_=maximal elongation velocity of individual sarcomeres. See **Fig. S7** for detailed data of each substance.

## Discussion

We introduce SarcAsM, a software tool for automated, rapid, and robust assessment of sarcomere structure and function in cardiomyocytes. Based on our custom deep learning strategy, SarcAsM analyzes more than 20 diverse and physically interpretable structural features, from individual Z-bands and sarcomere morphology to myofibril and cell-level sarcomere architecture, providing a comprehensive phenotyping of cardiomyocyte sarcomere morphology. In addition, SarcAsM tracks and analyzes the motion of individual sarcomeres in high-speed (66 frames per second) recordings with nanometer resolution (18 nm), providing important insight into dynamic phenomena such as sarcomere heterogeneity and “popping”^43^ that are hidden in traditional approaches that only determine whole-cell or average sarcomere dynamics.

The robustness and specificity of the analysis of local sarcomere lengths and orientations, resulting in the sarcomere vectors, builds on the specific and robust prediction of sarcomere Z-bands, M-bands and orientation using deep learning. Besides enabling precise Z-band morphometric analysis, we obtained a large set of sarcomere vectors, spatially describing sarcomere length. Furthermore the sarcomere vectors allow precise downstream analysis of higher-order structures as myofibrils and clustering of sarcomere vectors into domains, a measure for cardiomyocyte areas with presumably coordinated contractility. This function is particularly interesting in unconfined monolayer cultures with limited structural order.

Building on previous tools that each focus on distinct sarcomere features^22,23,26,29,44^, such as either Z-bands or sarcomeres, SarcAsM represents (1) a shift from traditional filter-based methods to context-aware structural deep learning and (2) a major methodological advancement by unifying previously disconnected analyses into a single comprehensive platform that spans all structural scales (from Z-band to cell level myofibril domains) with superior precision and efficiency. SarcApp^29^ pioneered the use of deep learning for Z-band segmentation and morphometrics, whereas SarcAsM extends this capability by simultaneously analyzing Z-bands, M-bands, and sarcomere orientation through its integrated deep learning framework. SarcTrack’s^22^ wavelet-based analysis of sarcomere lengths and orientations, while innovative, faces limitations in computational efficiency, resolution, and specificity that SarcAsM addresses through its more targeted deep learning-based sarcomere vector approach. Similarly, SarcGraph^26,30^, that analyzes sarcomere lengths based on Z-band segmentation and subsequent distance measurements, provides limited sampling of sarcomere structures and may lack precision when analyzing complex structures with elongated or non-linear Z-bands. In contrast, SarcAsM’s sarcomere vector methodology enables comprehensive sampling of all sarcomeres with enhanced precision and specificity.

SarcAsM is readily applicable to a wide range of microscopy data from muscle cells, including cardiomyocytes and skeletal muscle cells, with fluorescently labeled sarcomeres^11,22,45^ (**Fig. S2**). This versatility is possible because SarcAsM contains a pre-trained generalist neural network model for Z-band detection, trained on a diverse dataset encompassing different labeling methods and imaging modalities. SarcAsM robustly and specifically detects Z-bands even in challenging conditions, such as weakly fluorescent images or when Z-bands are distorted or obscured by artifacts or other highly fluorescent cellular structures. This capability not only helps avoid selection bias of regular over disfigured sarcomere morphology, but also enables effective discrimination between true sarcomeres and non-sarcomere fluorescent structures like stress fibers or cell adhesion sites.

We have developed SarcAsM as a user-friendly computational tool with broad applicability in cardiovascular research, for users with and without programming expertise. To achieve this, SarcAsM is available as standalone application with graphical user interface (**Fig. S8**) and as a well-documented Python package with a high-level API, that can be integrated into big data workflows. Due to its computational efficiency, SarcAsM runs on regular non-GPU computers for small and medium-sized datasets of hundreds of images (**Fig. S9**). On a GPU-equipped workstation or computing cluster SarcAsM can efficiently process terabyte-sized datasets of 10,000s of images.

SarcAsM is a versatile and precise computational tool suitable for a wide variety of studies of sarcomere structure and function, including morphogenesis and drug testing as well as detailed studies on sarcomere dynamics and biophysical studies of contractility^43^. Here we have demonstrated the utility of SarcAsM primarily in a genetically engineered ACTN2-citrine Z-band reporter model, but point out that SarcAsM can be applied to any sarcomere painting model. The application in the testing of acute and chronic drug activity in iPSC-derived cardiomyocytes should be of interest to the research and drug development community, as it may help to detect in preclinical animal models unrecognized unwanted effects. In this context, the finding of sarcomere disrupting effects of the actin-myosin inhibitor Mavacamten at clinically relevant concentrations (plasma concentrations of 350-700 ng/mL [1.3-2.6 µM]^38,46^) is of note, as it may offer a mechanistic explanation of side effects (such as worsening functional capacity, exacerbation of heart failure, and arrhythmia^47^), which are observed in the increasing number of Mavacamten-treated patients with hypertrophic cardiomyopathy. With the demonstration of the efficiency of SarcAsM in acute and chronic drug testing we anticipate its application in human-centric (hiPSC-based) high-content and high-throughput drug screens.

## Author contributions

DH, TD, CFS, and WHZ conceptualized the project; DH designed and wrote the software; DH and YL wrote the software documentation; AP built the graphical user interface; LH, TD, GB, BB, and MT performed experiments; LC created the ACTN2-Citrine iPSC-line; DH and LH annotated the training data; DH analyzed and visualized the data; DH drafted the manuscript; all authors reviewed and edited the manuscript.

## Acknowledgements

The authors gratefully acknowledge use of the microscopy facility of the Max Planck Institute for Multidisciplinary Sciences with access to high-speed confocal microscopy. We thank Laura Cyganek, Yvonne Hintz, Nadine Gotzmann, Lisa Schreiber, and Yvonne Wedekind (Stem Cell Unit, University Medical Center Göttingen) for excellent technical assistance. DH was supported by the German Academic Foundation (Studienstiftung des Deutschen Volkes) during his PhD studies and the Campus Institute for Data Science (CIDAS), University of Göttingen, for an early career postdoctoral fellowship. WHZ is supported by the DZHK (German Center for Cardiovascular Research), the German Federal Ministry of Education and Research (IndiHEART: 161L0250A, AutoOrgan_3R: LW23AT008), the German Research Foundation (DFG SFB 1002 C04/S01, IRTG 1816, RTG 2824, EXC 2067-1), and the Fondation Leducq (20CVD04). LC is supported by the German Research Foundation (DFG project number 417880571, project number 501985000, SFB 1002 S01, EXC 2067-1), and the Else Kröner Fresenius Foundation (project number 2019_A75). Generation of the line LiPSC-GR1.1 (also referred to as TC1133 or RUCDRi002-A; lot number 50-001-21) was supported by the NIH Common Fund Regenerative Medicine Program, and reported in Stem Cell Reports.^45^ The NIH Common Fund and the National Center for Advancing Translational Sciences (NCATS) are joint stewards of the LiPSC-GR1.1 resource. Repairon GmbH acquired and imported a vial of the TC1133 master cell bank, from which a Working Cell Bank (WCB) was created. myriamed GmbH acquired a derivative of the WBC from Repairon GmbH and provided a non-GMP derivative thereof to the Institute of Pharmacology and Toxicology at the University Medical Center Göttingen for non-commercial research use. DH and CS would like to thank the Isaac Newton Institute for Mathematical Sciences for support and hospitality during the program “New statistical physics in living matter: non-equilibrium states under adaptive control”.

## Additional information

Supplementary information is available for this paper at https://doi.org/xxxxx.

## Methods (online)

### Segmentation of Z-bands, M-bands, sarcomere orientation and cell mask using deep learning

SarcAsM employs a nested U-Net++^34^ convolutional neural network with three hierarchical levels and five output heads to jointly segment sarcomere sub-structures (Z-bands, M-bands), cell masks, and predict orientation fields. Classification tasks (Z-bands, M-bands, cell masks) use a combined binary cross-entropy and Dice loss to address class imbalance while preserving structural coherence. For the 2D sarcomere orientation field (unit vectors showing direction from M-band to Z-band) we use component-wise MSE+MAE losses with magnitude regularization to enforce unit consistency and adaptive masking of valid regions. The multi-task architecture enables shared feature refinement across classification and regression via unified encoder-decoder pathways.

As training data, we curated a diverse set of 220 images from various cell types, labeling methods, and microscopes - both our own and published datasets (refer to **Table S2** for details and sources). To create ground truth annotations, we employed a multi-step approach: For each image, the sarcomere Z-bands were manually traced using a graphics tablet, and binary masks were generated for cell regions. Subsequently, we leveraged a custom double-wavelet analysis, partly inspired by the SarcTrack^22^ algorithm (methodology described in **Supplementary Note 1, Fig. S10**), to create M-band masks, and sarcomere orientation fields. Briefly, we convolved images with a double-wavelet filter bank spanning different sarcomere lengths (1.4 to 2.7 µm at 0.02 µm intervals) and orientations (180° at 2° intervals). This computationally intensive process was accelerated using the GPU-accelerated PyTorch^48^ library. M-band masks were obtained by binarizing the maximum filter score above a threshold of 0.25 (max. score 1), corresponding to the midpoint between adjacent Z-bands. Sarcomere orientation fields were generated by drawing lines centered on M-band points (skeletonized M-band masks) with lengths and orientations corresponding to the local sarcomere lengths and orientations (corresponding to the maximum filter score) and the value of the orientation angle θ. This map was then smoothed using a median filter with a 3×3 kernel, excluding undefined regions. Prior to training, the angular orientation map was transformed into a 2D vector field (cos(*θ*), sin (*θ*)).

The training dataset was augmented sixfold through random rotations, Gaussian/Shot noise addition, and brightness/contrast adjustments to reflect diverse imaging conditions. This generalist pre-trained model allows most users to analyze their data without additional training. However, users can fine-tune or retrain models with custom datasets if necessary.

### Segmentation of Z-bands in high-speed time-lapse data

For high-speed time-lapse data of beating cardiomyocytes, with potentially low signal-to-noise ratios due to the fast imaging speed, a temporally consistent neural network architecture “3D U-Net”^35,36^ is used for Z-band segmentation, which treats a movie as 3D data (t, x, y) and allows more time-consistent segmentation by using both the spatial and temporal context of a signal (**Fig. S3a-c** and **Movie 1**). We adapted a small, parameter-efficient 3D U-Net architecture optimized for fast processing of microscopy movies^36^. As training data, 200 movie segments of 128 frames each were randomly selected from a diverse dataset of our ACTN2-citrine CM movies. Ground truth was created by a 2D U-Net and manually curated to remove flickering and short Z-band segments present in fewer than 4 frames. We augmented the training data by addition of noise and periodic drifts with random frequency and amplitude to specifically train rapidly moving Z-bands. Validation against the standard U-Net, trained with the same data, showed improvement in time-consistency (**Fig. S3d**). Time-consistency was quantified by the ratio of the sum of areas of Z-band objects present in more than 64 consecutive frames to the total area of Z-bands in 20 randomly selected movies of 128 frames each, which were distinct from the training data.

### Spatial analysis of sarcomere lengths and orientations by sarcomere ‘vectors’

Sarcomere vectors are constructed from the Z-band and M-band masks and the sarcomere orientation field is predicted by deep learning. The positions of the vectors are determined through a multi-step process: first, M-bands are thresholded and skeletonized to identify points along their 1-pixel thick boundaries; these points serve as the positions of the sarcomere vectors (M-points). The sarcomere orientation field is then transformed into an angle θ using the arctangent function (*θ* = arctan(*F*_*y*_/*F*_*x*_)), and its range is projected from 0°-360° to 0°-180°, accounting for the bipolar structure of sarcomeres. The resulting orientation angles are smoothed using a median filter with a 3×3 kernel. At each M-point, the local orientation angle (*θ*) is extracted as the sarcomere vector orientation. Finally, the sarcomere length is determined by analyzing the line intensity profile of the Z-band masks along the sarcomere vector orientation within a 3.5 µm length centered at the M-point. The Z-band peak positions are identified using the center-of-mass method, and the sarcomere length is calculated as the distance between the two innermost peaks when two peaks are found within a physiologically plausible range of 1-3 µm. If no such peaks are detected, the corresponding sarcomere vector is discarded. Details on parameters and implementation can be found in API documentation under `Structure.analyze_sarcomere_vectors`.

### Quantification of cell mask and area occupied by sarcomeres

The total area and area ratio occupied by cells are quantified for each image using the cell masks predicted by deep learning. Analogously, the total area of sarcomeres is calculated from the predicted sarcomere mask. The fraction of cell area occupied by sarcomeres is calculated by dividing the total area of sarcomere regions by the total area of cells. For more details on parameters and implementation, see the API documentation for `Structure.analyze_cell_mask`and `Structure.analyze_sarcomere_vectors`.

### Segmentation and morphometric analysis of Z-bands

To segment individual Z-bands, masks of Z-bands detected by deep learning are binarized by thresholding, and connected regions are uniquely labeled as objects. The length *L* of each Z-band object *j* is then calculated after removing objects shorter than a threshold length *L*_*min*_= 0.5 µm. To quantify the straightness of Z-bands, a straightness index *R* is calculated as the ratio of the object area to the convex hull object area (*R* = 1 is perfectly straight, *R ≪* 1 is curved). Additionally, the orientations of individual Z-bands *φ*_*j*_ are analyzed. On this basis, a global orientational order parameter (*OOP*) is computed for each cell. Since Z-bands are non-polar, the orientation angles *φ*_*j*_ range from 0° to 180°. *OOP* is a modification of the Kuramoto order parameter^49^ for bi-polar elements by mapping orientations from 0°-180° to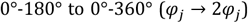: 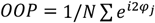where *N* is the number of Z-bands, *j* is the counting index and *i* is the imaginary unit.

*OOP* is <<1 for objects with random orientations between 0-180° ranging to 1 for uniaxially oriented Z-bands. The object labels also serve as masks to analyze the fluorescence intensity per area of each Z-band. At the cell level, the ratio of Z-band fluorescence to other fluorescence is evaluated. For more details on parameters and implementation, see the API documentation for `Structure.analyze_z_bands`.

### Lateral alignment and distance of sarcomere Z-bands

To analyze the lateral alignment and distance of Z-bands, masks of individual Z-band objects were skeletonized, and the endpoints of each object were determined using a specific filter (convolution with a 3×3 pixel kernel, returning 1 only if the center pixel and one of the border pixels are 1, otherwise 0). For each Z-band object, the positions, and orientations *Φ* of both ends are determined (**Fig. S4a**). Then, the pairwise Euclidean distances *D* between all Z-band ends are calculated. For all end pairs (*i, j*) in proximity (0.25 *μm < d <* 5 *μm*), the alignment index *A _i, j_* is defined as *a*_*i, j*_ for *a*_*i, j*_ ≥ *A*_*min*_, *and* 0 otherwise, with *a*_*i, j*_ = *cos*(*Φ*_*i*_ − *Φ*_*j*_+ *π*) · *cos*(*θ*_*i, j*_) · *cos*(*θ*_*j,i*_).

*A* is a custom metric based on the orientational difference of the Z-band ends |*Φ*_*i*_ − *Φ*_*j*_| as well as the offset between Z-bands, quantified by the orientational difference between the orientation of the Z-band *Φ* ends and the orientation of the vector connecting both ends *θ*_*j,i*_ (**Fig. S4a**). A is 1 for perfect antiparallel alignment of adjacent Z-bands and yields values smaller than 1 for Z-band pairs with offset or different orientations (**Fig. S4b**). A is 0 when the orientational difference between the two ends *Φ*_*i*_ − Φ_*j*_ is less than 90° and/or if one of the offset angles *θ*_*i, j*_ or *θ*_*j,i*_ is greater than 90°, i.e., if the Z-bands are interdigitating, and/or if it is smaller than *A*_*min*_ (**Fig. S4b**).

Finally, we used the Hungarian algorithm^50^ to solve the linear assignment problem, identifying the optimal set of paired Z-band ends with high alignment *A*_*i, j*_, linking each pair exactly once while maximizing the total alignment ∑*A*_*i, j*_. The resulting distributions of distances *D* and alignment indices *A* are an indicator of the higher-order lateral organization of Z-bands, with low distances and high alignment indices indicating more regular and ordered sarcomere structures. The sum of lengths of groups of laterally linked Z-bands *L*_2_, computed from a constructed graph, reflects the width of bundles of registered myofibrils, describing the effective width of cardiomyocytes. For more details on parameters and implementation, also see the API documentation for `Structure.analyze_z_bands`.

### Analysis of myofibril lengths by automated line growth algorithm

SarcAsM identifies myofibrils (uninterrupted series of aligned sarcomeres) using a custom line growth algorithm. Starting from randomly selected sarcomere vectors as seeds (typically 10% of all sarcomere vectors), lines grow bidirectionally following sarcomere orientations. For each growth step, the algorithm first estimates the next position based on the current sarcomere’s length and orientation, then refines this position using a next-neighbor approach. Growth continues until no suitable neighbors are found within a threshold distance (typically 0.4 µm). By calculating the estimate for the next step based on the average of the length and orientation of *p* adjacent segments of the line (e.g., *p* = 3), the persistence of the lines can be tuned, such that the growth algorithm becomes more robust to single outliers.

For each line, the length *L*, the global straightness *S*, calculated as *S* = 1 *− d_max_*/*d*_*ends*_ with the end-to-end distance *d*_*ends*_, the maximal projected distance of the line to the end-to-end line *d*_*max*_, and the bending energy *B* = 1/*N* ∑(*θ*_*i*+1_ − *θ*_*i*_)^2^, as a measure for the local wiggliness of the myofibrils, are calculated. For each image, the distribution mean, and standard deviation of line lengths and MSCs are calculated, serving as estimators for myofibril order and variability. For more details on parameters and implementation, see the API documentation for `Structure.analyze_myofibrils`.

### Clustering of myofibril domains

SarcAsM detects domains of interconnected sarcomeres with similar orientation. For that purpose, a graph representing the network of sarcomere vectors is constructed: sarcomere vectors are nodes and edges are added between sarcomere vectors if the ends are within a distance threshold of, e.g., 0.5 µm with the absolute cosine similarity as weight (*w*_*i,j*_ = | cos(*θ*_*i*_) − cos(*θ*_*j*_)| for vector pair *i, j*), penalizing orientational differences. Edges with *w*_*i,j*_ < 0.6, for example, are removed.

Subsequently, the Leiden graph-based community detection algorithm^37^ is applied to cluster spatially connected and similarly oriented sarcomere vectors into domains. The resolution parameter, set to, e.g., 0.05, controls the granularity of the detected clusters. Higher resolution values lead to detecting smaller, more fine-grained clusters, while lower values result in larger, more coarse-grained clusters.

The area of each domain is quantified based on its sarcomere vectors, and domains with areas smaller than, e.g., 50 µm^2^ are omitted. In each image, the number of domains and their areas are quantified, as well as the mean and standard deviation (STD) of sarcomere lengths and the orientational order parameter (OOP) of the sarcomere orientations in each domain. For more details on parameters and implementation, see the API documentation for `Structure.analyze_sarcomere_domains`.

### Automated or manual LOI selection for sarcomere motion tracking

SarcAsM offers two approaches for identifying suitable lines of interest (LOIs) for tracking sarcomere dynamics: manual selection via the graphical user interface (GUI) or automated detection based on local sarcomere order, the latter being particularly valuable for processing large datasets.

For automated LOI detection, the myofibril line growth algorithm is applied with a large persistence parameter (e.g., p=5) to generate a set of lines (using, e.g., 10 % of sarcomere vectors as seeds). This set is then filtered to retain lines with more than, e.g., 10 sarcomeres. Additional filtering based on straightness, sarcomere length statistics, or midline lengths can select regions with evenly spaced sarcomeres.

To obtain distinct LOIs from different regions, similar lines are grouped using a pairwise Hausdorff distance metric as a similarity measure. Agglomerative clustering based on these distances identifies similar lines. From each cluster, either the longest line or a random line is selected as an LOI (allowing curved LOIs) or a straight line is fitted to the cluster points with length constrained to the point cloud size.

Along each LOI, intensity profiles *I*(*x, t*) are extracted from the Z-band movie detected by deep learning. The line width (here 10 pixels ∼0.72 µm) averages intensity perpendicular to the line direction for a more robust signal. For parameters and implementation details, see the API documentation for `Structure.detect_lois`.

### Detection and tracking of individual Z-band and sarcomere motion

Individual Z-band positions are determined from the intensity profiles *I*(*x, t*) of each LOI. For each time point, the position of each Z-band *i* is calculated through a two-step procedure: first identifying the intensity maximum as an initial approximation, then determining the exact center *z*_*i*_ using the center-of-mass method.

SarcAsM links peak positions *z*_(*i,t*)_ across all frames to create continuous Z-band trajectories *Z*_*i*_(*t*) using the Crocker-Grier algorithm^41^ for particle tracking. This algorithm associates peaks over a memory time interval, improving tracking robustness. Z-band tracking precision was estimated to be ∼17 nm by determining the standard deviation of *Z*_*i*_(*t*) around its mean position 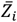 in quiescent (non-contracting) cardiomyocytes (**Fig. S3e-g**).

Z-band trajectories *Z*_*+*_(*t*) are smoothed using a Savitzky-Golay filter (default parameters: window length 11, polynomial order 5) to remove high-frequency fluctuations and isolated outliers. Individual sarcomere length time series *SL*_*i*_ (*t*) are calculated as the distance between neighboring Z-bands: *SL*_*i*_(*t*) = *Z*_*i*+ 1_(*t*) − *Z*_*i*_(*t*). Sarcomere velocity time series *V*_*i*_ (*t*) are calculated by temporally differentiating sarcomere lengths: *V*_*i*_ (*t*) = (*SL*_*i*_ (*t* + 1) − *SL*_*i*_ (*t* − 1))/(2Δ*t*), where Δ*t* is the frame time. For each LOI, average sarcomere length 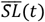 and average velocity 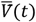 are also calculated.

For implementation details and additional parameters, see the API documentation for `Motion.detect_peaks`and `Motion.track_z_bands`.

### Detection of sarcomere contraction intervals using deep learning

The contraction state (*S*(*t*)) (0 = quiescent, 1 = contraction) and the start and end time points of each contraction interval are detected using a custom deep learning approach. Our ContractionNet model detects contraction intervals from time-series data of individual Z-band positions and sarcomere lengths (**Fig. S6a**). The model architecture consists of two initial convolutional layers (kernel size 5), followed by a dilated convolution layer capturing broader temporal patterns. Each convolution includes instance normalization and ReLU activation. A self-attention layer enhances focus on salient features amid noise before two final convolutional layers and a sigmoid activation function producing binary output S(t). The model was trained on 50 manually annotated LOIs plus simulated periodic data with various frequencies, amplitudes, and noise levels. Data augmentation included Gaussian fluctuations, random outliers, and baseline drifts. Training ran for 100 epochs using a combined BCE-Dice loss (learning rate 0.001).

For each LOI, individual sarcomere and Z-band predictions *S*_+_(*t*) are combined into a cell-level contraction state: *S*(*t*) = 1 *if* ⟨*S*_*i*_(*t*)⟩_*i*_ ≥ 0.5, *else S*(*t*) = 0. From *S*(*t*), we calculate spontaneous beating rate (BR), beating rate variability, contraction duration (T_C), and quiescent phase duration (T_Q) for each LOI. For implementation details, see the API documentation for `Motion.detect_peaks`and `Motion.detect_analyze_contractions`.

ContractionNet significantly outperformed threshold-based approaches on validation data (median F1-score 0.96 vs. 0.81) and maintained excellent performance (F1 score > 0.9) on simulated data with signal-to-noise ratios down to 3.4 dB (**Fig. S6b-g**).

### Analysis of individual and average sarcomere trajectories

SarcAsM quantifies sarcomere dynamics by calculating equilibrium lengths (*SL*_0,*i*_) during quiescent phases (*S* = 0) and measuring length changes (Δ*SL*_*i*_(*t*) = *SL*_*i*_ (*t*) − *SL*_0,*i*_) relative to this baseline. For each contraction cycle j, the software assesses extrema of sarcomere length changes (maximal shortening and lengthening amplitudes) for each individual sarcomere *i* (ΔSL_−,*i,j*_, ΔSL_+*i,j*_) and multi-sarcomere averages 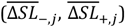. Similarly, velocity extrema are quantified at both individual (*V*_−,*i,j*_, *V*_+*i,j*_) and averaged 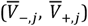 levels. Temporal parameters include time to peak shortening (T_p_) and custom contraction kinetics (e.g., time from 10% to 90% shortening), providing comprehensive characterization of sarcomere contractile behavior. For detailed implementation specifics and additional parameters, see the API documentation for `Motion.detect_analyze_contractions`.

### Algorithm runtimes and hardware requirements

We assessed algorithm runtimes on an Nvidia H-100 GPU within an HPC cluster, a GPU-equipped workstation (Windows 10, Nvidia Quadro RTX 5000), and a notebook computer (Ubuntu, CPU: Intel Core i7-10750H) (**Fig. S9**). While large datasets were analyzed on the H-100 for optimal performance, SarcAsM is optimized for efficiency and can be run on workstations or non-GPU computers for small to medium-sized datasets (∼100s of images). For example, analyzing a single 2,000×2,000-pixel image takes around 10 seconds on a standard workstation.

Models were trained for 200 epochs at a learning rate of 0.0001 using the Adam optimizer on the H-100 GPU. While the H-100 offers faster training for large datasets, training is also feasible on GPUs like the Nvidia Quadro RTX 5000, albeit with longer runtimes. SarcAsM provides a generalist pre-trained model suitable for most users, but custom training or fine-tuning may be required for specialized datasets and custom high-speed data. Instructions for creating custom datasets and training models are available in the online documentation (https://sarcasm.readthedocs.io/en/).

### Validation of SarcAsM

SarcAsM’s performance was evaluated focusing on two key areas: (1) detecting sarcomeres and quantifying sarcomere lengths in individual images, and (2) dynamically tracking individual and average sarcomere motion in high-speed recordings of living cardiomyocytes. A diverse set of 10 images was curated, encompassing various sarcomere labeling methods, imaging modalities, cell types, and experimental conditions. To specifically test the tool’s cross-applicability, this validation set included several images representing data types completely absent from the training dataset.

Sarcomere detection was evaluated using precision, recall, and F1 score against manually annotated ground truth masks created with napari^51^. We defined sarcomeres as at least two parallel adjacent Z-bands, each with clear linear character, not dots. To validate sarcomere length analysis, we manually measured sarcomere lengths at 100-200 positions across each image as ground truth. For motion tracking, the continuity, noise level, and physiological plausibility of individual and average sarcomere trajectories was evaluated. Detailed results of this benchmarking are presented as **Supplementary Note 2**. All samples and code with analysis parameters are available as **Supplementary Materials**.

### Generation of hiPSC ACTN2-Citrine reporter line

Derivatives of the hiPSC line TC1133^45^, which was developed under informed consent (refer to https://hpscreg.eu/cell-line/RUCDRi002-A), were used to engineer the ACTN2-Citrine reporter line. Genomic integration of citrine coding sequence at the C-terminal 3’-end of the last exon of the ACTN2 gene was performed using CRISPR/Cas9 technology (**Table S1**). The guide RNA target sequence was 5’-ATCGCTCTCCCCGTAGAGTG-3’. The donor vector consisted of the citrine coding sequence (717 bp) flanked by left and right homology arms (680 bp each). To inhibit cleavage of the donor strand, several point mutations were introduced in the sequence complementary to guide RNA. In addition, an 18 bp sequence encoding 5(Gly)-Ala was inserted at the 5’ end of the citrine cDNA to minimize potential folding perturbations between ACTN2 and citrine. The donor DNA was cloned into pUC57 (GENEWIZ, Takeley, UK). For editing, the CRISPR/Cas9 ribonucleoprotein complex was assembled by mixing of the Alt-R CRISPR-Cas9 crRNA and the Alt-R CRISPR-Cas9 tracrRNA (preassembled in a 1:1 ratio) with the Alt-R Hifi SpCas9 Nuclease 3NLS (IDT DNA Technologies) at 1:3 molar ratio together with the donor plasmid for homology-directed repair in nucleofector solution. Transfection with premixed CRISPR/Cas9 solution was performed with the 4D Amaxa Nucleofector system (Lonza; program CA-137) and the P3 Primary Cell 4D-Nucleofector X Kit (Lonza) according to the manufacturer’s instruction. Following nucleofection, hiPSCs were replated into a Matrigel-coated well of a 6-well plate containing StemFlex medium supplemented with 2 µM Thiazovivin and 100 U/ml penicillin and 100 µg/ml streptomycin (Thermo Fisher Scientific). After 3 days, transfected hiPSCs were singularized using the single cell dispenser CellenOne (Cellenion/Scienion) in StemFlex medium on Matrigel™-coated 96-well plates. Successful genome editing was identified by PCR and Sanger sequencing using the primers 5’ – GGAATTGTCCTATTTCCCACTG-3’ (forward) and 5’-GCATGAAAATAAAACATTAGAATCC - 3’ (reverse). The heterozygously tagged hiPSC line RUCDRi002-A-3 clone 51J5/TC-1133-ACTN2-Citrine (in short ACTN2-citrine) was expanded and maintained in StemMACS iPS-Brew XF medium on Matrigel-coated plates.

### Protein electrophoresis and immunoblotting

500,000 differentiated cardiomyocytes were lysed in 150 µl lysis buffer, containing 150 mM NaCl, 50 mM Tris at pH 7.4, 2 mM EDTA and 1% Nonidet P-40. Then, 6x Laemmli loading buffer was added and the sample was heated to 95°C for five minutes for protein denaturation. 15 µl of the sample was resolved by a 12% NuPAGE Bis-Tris gel (Invitrogen) and proteins were transferred onto a polyvinylidene difluoride (PVDF) membrane (Immobilon-P; Millipore, Billerica, MA) using a Bio-Rad Criterion Blotter. After protein transfer, the membrane was blocked with 10% non-fat milk in PBS containing 0.05% for 20-30 minutes and incubated with affinity-purified monoclonal anti-ACTN2 antibody from mouse (A7811, Sigma-Aldrich) diluted at 1:2,500 or anti-GFP antibody (11814460801; Roche) diluted at 1:500 at 4 °C overnight, followed by treatment with 1:5,000 diluted peroxidase-labeled goat anti-mouse IgG (A4416, Sigma-Aldrich). Reactivity was detected with an Amersham ECL Detection Reagent (GE Healthcare Life science) using an Intas gel imager (Intas, Göttingen, Germany). The membrane probed with anti-ACTN2 antibodies was stripped with buffer containing 2% SDS, 100 mM β-Mercaptoethanol, and 62.5 mM Tris-HCl (pH 6.8) at 50°C for 30 min to remove bound probes and re-probed with 1:50,000 monoclonal anti-GAPDH antibodies from mouse (60004-1-Ig, proteintech) in combination with 1:10,000 peroxidase-labeled goat anti-mouse IgG (A4416, Sigma-Aldrich).

### Genotyping and sequencing

500,000 cardiomyocytes were lysed in 50 µl of lysis buffer, containing 50 mM KCl, 10 mM Tris at pH 8.3, 2.5 mM MgCl2, 0.45% Nonidet P-40, 0.45% Tween 20, 0.01% gelatin and 1 µg/mL Proteinase K at 65 °C for 30 min. Proteinase K was then inactivated by incubation at 95 °C for 15 minutes and a 10 µL sample was used as template for PCR. Primers for DNA amplification were 5’-GGAATTGTCCTATTTCCCACTG-3’ (forward) and 5’-GCATGAAAATAAAACATTAGAATCC-3’ (reverse). PCR products were cloned into pCR® 2.1vector using TA cloning® kit (Invitrogen) and sequenced by Seqlab (Göttingen, Germany) using M13 primers.

### Cardiomyocyte differentiation and culture

Cardiomyocyte differentiation of hiPSCs was performed as described in Tiburcy *et al*.^10^. hiPSC-ACTN2-Citrine-derived cardiomyocytes were cultured in 6-well plates (Cat# 3516, Corning) in serum-free “cardio” medium (0.4 mM Ca2+, RPMI 1640 with GlutaMAX (Cat 61870, Invitrogen), 1% penicillin/streptomycin (Cat 15140, Invitrogen), 2% B27 supplement (Cat 17504-044, Invitrogen) at 37°C in a 5% CO_2_ incubator with culture medium changes every other day. For re-seeding purposes, cells were detached using Accutase^®^ digestion medium (StemPro® Accutase® cell dissociation reagent (Cat A11105-01, Gibco), 0.025% Trypsin (Cat 15090-046, Gibco), 20µg/mL DnaseI (Cat 260913, Calbiochem) for 15-20 min at 37°C. Digestion was stopped using “cardio” medium supplemented with 5 µM Rock Inhibitor (Stemolecule Y27632, Cat 04-0012-10, Reprocell) at three times the amount of Accutase^®^ mix. Cell clumps were separated using a 100 µm cell strainer and cells were cultured 24 hours with 5 µM Rock Inhibitor. Cells were seeded at approximately 150,000 cells per micropatterned substrate and imaged 20-30 days after seeding.

### Fabrication of cell-adhesive micropatterns on polyacrylamide soft gels

Photoresist masters for polydimethylsiloxane (PDMS) stamps were fabricated via soft lithography in a Class 100 cleanroom. Silicon wafers (Microchemicals GmbH, Germany) were spin-coated with 2 mL of SU-8 3005 negative photoresist (MicroChem, USA) at 500 rpm for 10 s (ramp rate: 100 rpm/s), followed by 1,800 rpm for 30 s (ramp rate: 300 rpm/s). After a soft-bake at 95°C for 5 min, wafers were exposed to UV light (366 nm) using custom soda-lime photomasks (Compugraphics, Germany) for 5 s. Post-baking was performed at 65°C for 1 min and 95°C for 3 min. Non-illuminated regions were developed in AV-rev 600 developer, and the masters were hard-baked at 150°C for 5 min, yielding an 8 µm thick photoresist pattern.

PDMS stamps were prepared using Sylgard® 184 PDMS and curing agent (Dow Corning, USA) mixed at a 10:1 ratio. The mixture was degassed under vacuum for 20 min, poured onto the photoresist masters, and degassed again for 15 min. After curing at 60°C for 2 hours, the PDMS was peeled off and cut into ∼1 cm × 1 cm octagonal stamps. Stamps were plasma-cleaned for hydrophilicity (5 min) and incubated overnight with 150 µL of Synthemax™ II-SC coating material (Corning®, USA; Cat#3535) dissolved in ultrapure water. After wicking off the protein solution and air-drying, the stamps were transferred onto plasma-cleaned glass coverslips under a 50 g weight for 30 min to transfer the coating.

For gel preparation, acrylamide (Sigma-Aldrich), bis-acrylamide crosslinker, and PBS were mixed as a stock solution. Polymerization was initiated by adding ammonium persulfate (APS; Vizag Chemicals, India) at a ratio of 1:100 and tetramethylethylenediamine (TEMED; Muby Chemicals, India) at a ratio of 1:1,000. The gel modulus was adjusted to 15 kPa and confirmed by rheometry (Physica MCR 501 rheometer; Anton Paar, Austria). A total of 35 µL of gel solution was pipetted onto glutaraldehyde-coated glass slides (0.5% glutaraldehyde in ultrapure water), sandwiched with Synthemax™-patterned coverslips, and polymerized for 1 hour. The patterned gels were stored in PBS until use. One day before experiments, the top coverslip was removed gently, and gels were washed three times in PBS.

### Chronic and acute drug treatment

For chronic drug treatment, cells were seeded in 96-well plates (Cat 220.230.711, zell-kontakt) coated with Matrigel (1:120 diluted in 1X PBS; BD Biosciences, 354320) at 10,000 cells per well and imaged 1-8 days after seeding (9 fields per well, 4 h imaging interval, 7 z-slices with 1 µm spacing, z-slice with maximum intensity was analyzed). Drug exposure with Mavacamten (MYK-461, Cat S8861, Selleckchem) was started on day 3 and refreshed with daily medium changes. We computed a multivariate extension of the classical Z-factor^52^ by calculating the Euclidean distance between mean feature vectors of positive and negative controls relative to their average feature-wise standard deviations.

For acute drug treatment, ACTN2-citrine cardiomyocytes were cultured on micropatterned polyacrylamide gels with 15 kPa Young’s modulus. After 20-30 days of maturation, cardiomyocytes were treated with Isoprenaline (I5627, isoprenaline hydrochloride, Sigma Aldrich), Mavacamten (MYK-461, Cat S8861, Selleckchem), Levosimendan (Cat L5545, Sigma Aldrich), Omecamtiv mecarbil (CK-1827452, Cat S2623, Selleckchem), Verapamil (CP-16533-1, Cat S4202, Selleckchem), Digitoxin (Cat D5878, Sigma Aldrich), each dissolved in DMSO (10 mM stock solutions), with the final DMSO content in the experiments not exceeding 0.1%. After an incubation period of 3-5 min, spontaneously beating cells were manually selected and recorded up to 30 min after addition of the respective drug. A total of 1,804 cells were recorded. 6,944 LOIs were automatically identified, of which 3,594 LOIs in 1,129 cardiomyocytes met our quality criteria (*≥*10 sarcomeres, *≥*10 contraction cycles, aperiodicity < 0.25) and were included in the analysis.

### Microscopy

High-speed live-cell microscopy of micropatterned polyacrylamide gels was performed on a Leica SP5 scanning confocal microscope (TCS SP5 II, Leica, Germany) at 37°C and 5% CO2, using an 100x/1.4 NA oil immersion objective. Substrates were mounted in a custom-fabricated chamber for round Ø25 mm glass cover slides and kept in culture medium during imaging. We used an 8 kHz resonant scanner with bidirectional scanning mode and recorded up to 20-30 s movies with 1,024 × 200 pixels and a temporal resolution of 66 frames per second (example see **Movie 1**). Long-term live cell microscopy of 2D cardiomyocytes for drug testing was performed on a Yokogawa CV8000 spinning-disk confocal microscope (CellVoyager CV8000, Yokogawa, Germany) at 37°C and 5% CO2, using a 60x/1.2 NA water-immersion objective (example see **Movie 2**).

## Data availability

All training data, test data, validation data and exemplary data of chronic and acute drug screening experiments are available on Zenodo (https://doi.org/10.5281/zenodo.8232838).

## Code availability

SarcAsM is implemented in Python and uses the following secondary packages: NumPy and SciPy^53,54^ for array programming and scientific computing, Scikit-image^55^ for image processing and morphometric analysis, Scikit-learn^56^ for clustering, PyTorch^48^ for deep learning and Napari^51^ as image viewer. For 3D U-Net and regular U-Net, our own custom implementation is used (https://github.com/danihae/bio-image-unet). SarcAsM is available on GitHub (https://github.com/danihae/SarcAsM, public upon publishing) and Python Package Index (PyPI; https://pypi.org/project/sarc-asm/).

SarcAsM features high-level functions for exporting and plotting data. SarcAsM has detailed online documentation with instructions for installation and API usage (https://sarcasm.readthedocs.io/en/). The repository contains Jupyter notebooks with documentation and tutorials. SarcAsM is further available as pre-packaged application with a graphical user interface (GUI) comprising all main functions, visualizations, and batch processing (**Fig. S8**). For optimal performance, SarcAsM requires sufficient sampling of sarcomere structures; we recommend using images with pixel sizes of <0.2 μm. For support and questions, please contact us directly or use the issue tracking feature on our GitHub repository.

