## Supplementary Information for "SarcAsM: AI-based multiscale analysis of sarcomere organization and contractility in cardiomyocytes"

**Supplementary Information (SI) Guide:**


**Supplementary Note**

Supplementary Note 1 – **Generation of training data for AI-based sarcomere analysis**

Supplementary Note 2 – **Validation of sarcomere detection and length measurement**

**Supplementary Data**

Links provided for data transparency, reproducibility, and benchmarking.

**Supplementary Figures**

Supplementary Figure 1 - **Characterization of ACTN2-citrine Z-band reporter model**.

Supplementary Figure 2 - **Validation of 3D U-Net for Z-band segmentation and tracking.**

Supplementary Figure 3 - **Z-band lateral connections and sarcomere domain analysis.**

Supplementary Figure 4 - **Training data generation**.

Supplementary Figure 5 - **Automated line-of-interest detection for motion tracking.**

Supplementary Figure 6 - **Network architecture and validation of ContractionNet.**

Supplementary Figure 7 - **Acute drug effects on sarcomere dynamics**.

Supplementary Figure 8 - **Application of SarcAsM to different data sets**.

Supplementary Figure 9 - **SarcAsM graphical user interface of standalone application.**

Supplementary Figure 10 - **SarcAsM run times for different representative data.**

**Supplementary Table**

Supplementary Table 1 - **Schematic representation of ACTN2 edited alleles.**

Supplementary Table 2 - **Details and sources of training data for sarcomere model analysis.**

**Supplementary Movies**

Supplementary Movie 1 - **Detection of Z-bands from confocal microscopy movie.**

Supplementary Movie 2 - **Time-course of chronic drug treatment of ACTN2-citrine CMs.**

Supplementary Movie 3 - **Animation of sarcomere motion tracking workflow.**

Supplementary Movie 4 - **Analysis of simulated microscopy movie.**

**SI – Supplementary Note 1**

**Generation of training data for AI-based sarcomere analysis**

We created training data for sarcomere feature detection using a multi-step approach. Initially, manual tracing of sarcomere Z-bands was performed on each image in our diverse training dataset. A U-Net convolutional neural network[1] was then trained to predict sarcomere Z-bands, with extensive data augmentation applied to the training images to enhance model robustness. This augmentation included random variations in contrast, brightness, noise, rotation, and other parameters, allowing the U-Net to learn invariant features and generalize well to diverse input data. Utilizing the U-Net output, rather than the hand-drawn traces, for further processing is preferable because it provides a more accurate and generalized representation of the data. The U-Net predictions can mitigate potential inaccuracies in manual annotations, resulting in more robust training data for the subsequent processing steps.

Since sarcomeres are bounded by two adjacent Z-bands, it is possible to filter specifically for sarcomere structures by convolving the predicted Z-band mask (**Fig. S10a(1)**) with a double wavelet filter[2]. SarcAsM employs a filter bank of double wavelet filters (univariate Gaussians in the longitudinal direction limited by step function in the lateral direction) for a range of wavelet distances *SL* and orientations *θ*:

$$W^{\pm}(SL,\theta)_{x,y} = \frac{1}{\sqrt{2\pi}\sigma} exp\left(-\frac{(x'(\theta) \pm SL/2)^2}{2\sigma^2}\right) \times \begin{cases} 1, & |y'(\theta)| \leq w \\ 0, & |y'(\theta)| > w \end{cases}$$

with $x' = x\,cos(\theta) - y\,sin(\theta)$ and $y' = x\,sin(\theta) + y\,cos(\theta)$. The parameters $\sigma$ and $w$ set the widths of the wavelets (e.g. $\sigma = 0.1$ µ$m$, $w = 1$ µ$m$). The width $\sigma$ needs to be adjusted depending on imaging modality and sample, e.g., magnification or image quality. We used a range of sarcomere length from 1.4 to 2.6 µm with 0.025 µm steps, and orientation steps of 10°.

The two Gaussian filter banks $W^{\pm}(SL,\theta)_{x,y}$ were applied separately to the image of Z-bands predicted by deep learning, and both results were then multiplied pixel-wise with ($*$ = convolution, · = pixel-wise multiplication):

$$Z(SL,\theta)_{(x,y)} = \left(I_{(x,y)} * W^{+}(SL,\theta)_{(x,y)}\right) \cdot \left(I_{(x,y)} * W^{-}(SL,\theta)_{(x,y)}\right)$$

This is equivalent to an AND logic gate, yielding only positive results if both filter results are positive, i.e., sarcomere Z-bands are on both sides. The resulting 4-dimensional matrix $Z(SL,\theta)_{(x,y)}$ contains the result for each filter in the bank. The best-fitting filter, i.e., the filter with the highest score $Z$, reflects the local sarcomere length $SL$ and orientation $\theta$. To obtain $SL$ and $\theta$ locally, the filter with the maximal score $Z^{max}_{(x,y)}$ is determined at each pixel $(x,y)$ and the corresponding $SL$ and $\theta$ are stored as spatial maps:

$$\left(SL_{(x,y)},\theta_{(x,y)}\right) = argmax_{(SL',\theta')\in W}\left(Z(SL',\theta')_{(x,y)}\right)$$

A binary mask of regions with high scores $Z^{max}_{(x,y)}$, located at the sarcomere M-bands between parallel Z-bands, are created by binary thresholding of $Z^{max}_{(x,y)}$ with a threshold of, e.g., 0.25 of the maximum of $Z^{max}_{(x,y)}$ in an image or data set. The identified regions in the center between Z-bands serve as preliminary M-band mask (**Fig. S10a(3)**). Next, the M-bands are skeletonized, and the images of skeletonized lines are transformed into a set of M-point locations $\{(x_i, y_i)\}$. For each element of this set the values of $SL_{(x,y)}$ and $\theta_{(x,y)}$ are evaluated at location $(x_i, y_i)$, resulting in a set of sarcomere vectors $\{S_i\}$ with $S_i = (x_i, y_i, SL_i, \theta_i)$ (**Fig. S10a(4)**).

The set of sarcomere vectors is transformed into a continuous binary mask of sarcomeres (1 = sarcomere, 0 = no sarcomeres), by drawing binary lines and subsequent smoothing (**Fig. S10a(5)**). Similarly, the vectors are transformed into a continuous map of sarcomere orientation angles by drawing centered lines centered at the location of each sarcomere vector with values representing the vector angle from the M-band towards the closest Z-band. Undefined regions are set to NaN, and the angle map is smoothed using a median filter with radius of, e.g., 3 pixels, that ignores NaN values. To avoid artifacts caused by the angular discontinuity at 2π → 0, we transformed the angular map into a 2D unit vector field $\vec{V} = (\cos(\theta), \sin(\theta))$ prior to filtering, and then transformed it back by $\theta = \arctan\left(\frac{V_y}{V_x}\right)$, facilitating training data augmentation (**Fig. S4a(6)**).

The resulting ground truth contains a considerable number of false-positive sarcomere vectors, between neighboring myofibrils or when Z-bands are fragmented (**Fig. 10a(4)**). To address this issue, we manually corrected the M-band masks to remove false-positive sarcomere vectors, resulting in clean and precise ground truth for M-bands, sarcomere orientation, and sarcomere mask (**Fig. S10b(3,4,5,6)**). For further details on the training data generation process, including parameter settings and implementation, please refer to the API documentation for the class `TrainingDataGenerator` and the accompanying tutorial notebook, which provides an interactive environment for creating custom training data.

**SI – Supplementary Note 2**

**Validation of sarcomere detection and length measurements**

This note details the quantitative validation of SarcAsM's performance in segmenting sarcomere-containing regions (generating sarcomere masks) and measuring sarcomere lengths.

To rigorously assess accuracy and generalizability, we used a dedicated test set of 10 diverse images, distinct from the SarcAsM training data. This set included images from our own experiments (**Samples 1, 3, 6, 9, 10**) and external sources (**Samples 2, 4, 5, 7, 8**). Notably, several images (**Samples 4, 5, 7, 8**) represented data types completely absent from the training set, allowing for a robust evaluation of cross-applicability across different imaging modalities and conditions.

Validation compared SarcAsM's outputs against manually generated ground truth. Segmentation performance (sarcomere mask) was evaluated using F1-score, precision, and recall. Length measurement accuracy was assessed by comparing SarcAsM's results to extensive manual measurements within each image (details see main **Methods**, results in Figure below).

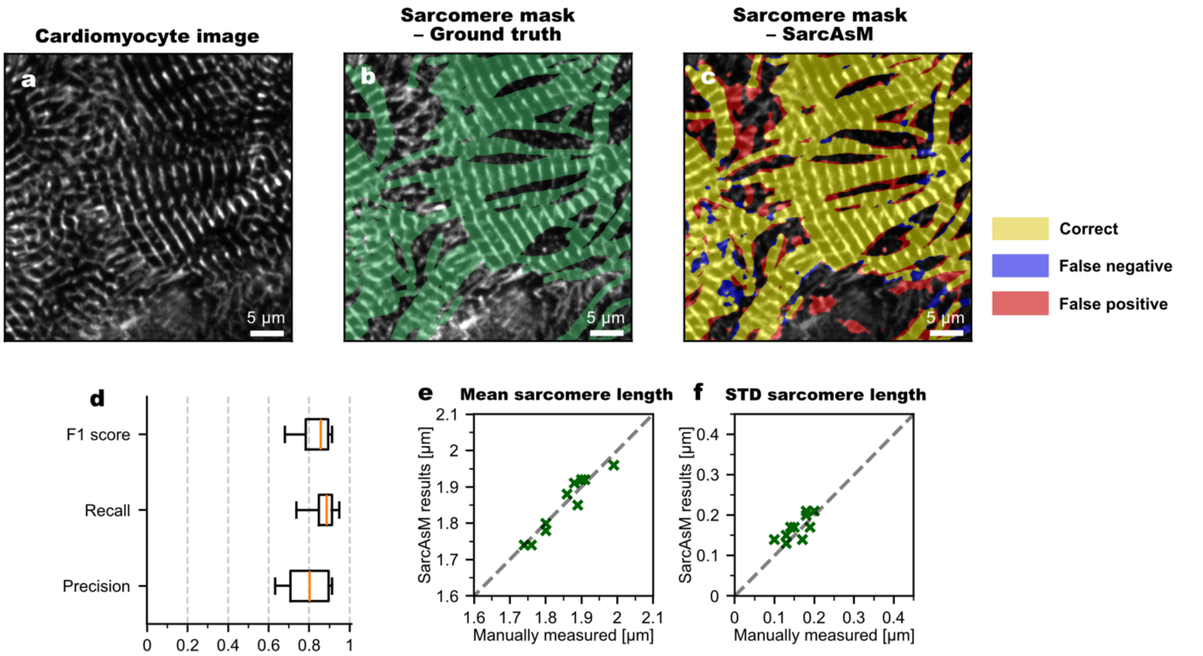


**Validation of SarcAsM's sarcomere detection and sarcomere length measurements.** (**a**) Representative image of ACTN2-citrine cardiomyocyte. (**b**) Manually masked regions with sarcomeres (green) as ground truth. (**c**) False color representation of sarcomere mask detected by SarcAsM validated against ground truth. Yellow regions mark correctly identified sarcomeres; blue regions highlight false-negative and red regions false-positive sarcomere detections. (**d**) Segmentation performance metrics for sarcomere detection. Precision: proportion of detected sarcomeres that are true positives; Recall: proportion of actual sarcomeres correctly identified; F1-score: harmonic mean of precision and recall. Boxplot shows quartiles (boxes), data range (whiskers) and medians (red lines). (**e,f**) Validation of sarcomere length analysis of manually measured (measured at >100 positions in each image using the line lengths tool in Fiji) against automatically measured sarcomere lengths with SarcAsM. Scatter plots show mean and standard deviation of distributions of manually measured against SarcAsM measurements in each image (N=10 diverse images, results see below **Samples 1-10**).

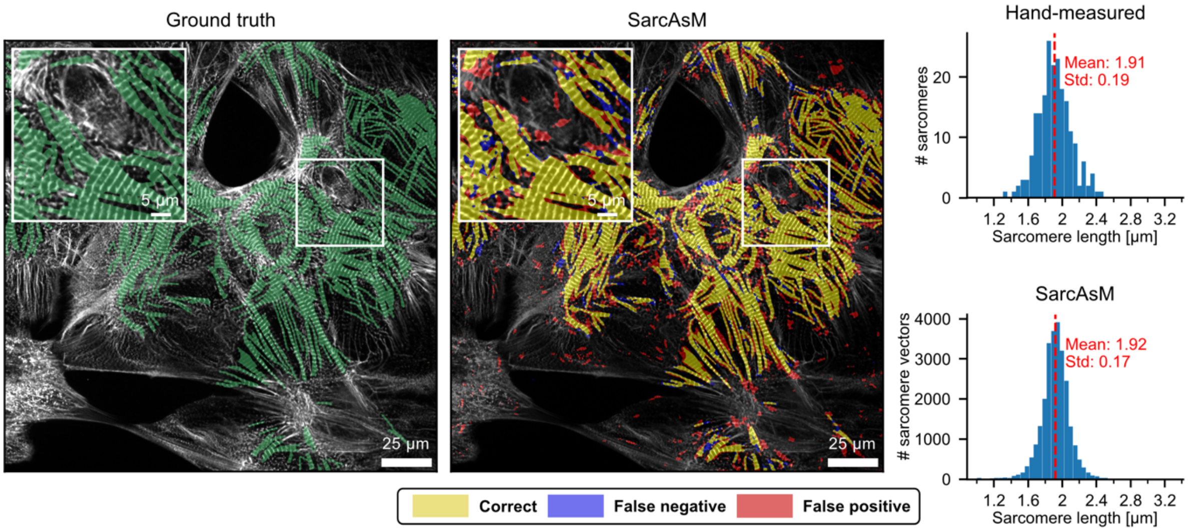


**Sample 1: hiPSC-derived ACTN2-citrine cardiomyocytes in 2D culture (own data). Left panel**: Microscopy image with green overlay depicting the ground truth sarcomere mask. **Middle panel**: SarcAsM algorithm output displayed as a false color map of sarcomere predictions, where yellow indicates correct predictions, blue represents false negatives, and red shows false positives. Segmentation performance metrics: precision 0.69, recall 0.89, F1-score 0.78. **Right panel**: Comparative histograms of sarcomere length distributions obtained through manual hand-measurement and analysis with SarcAsM. Respective means and standard deviations are shown in red.

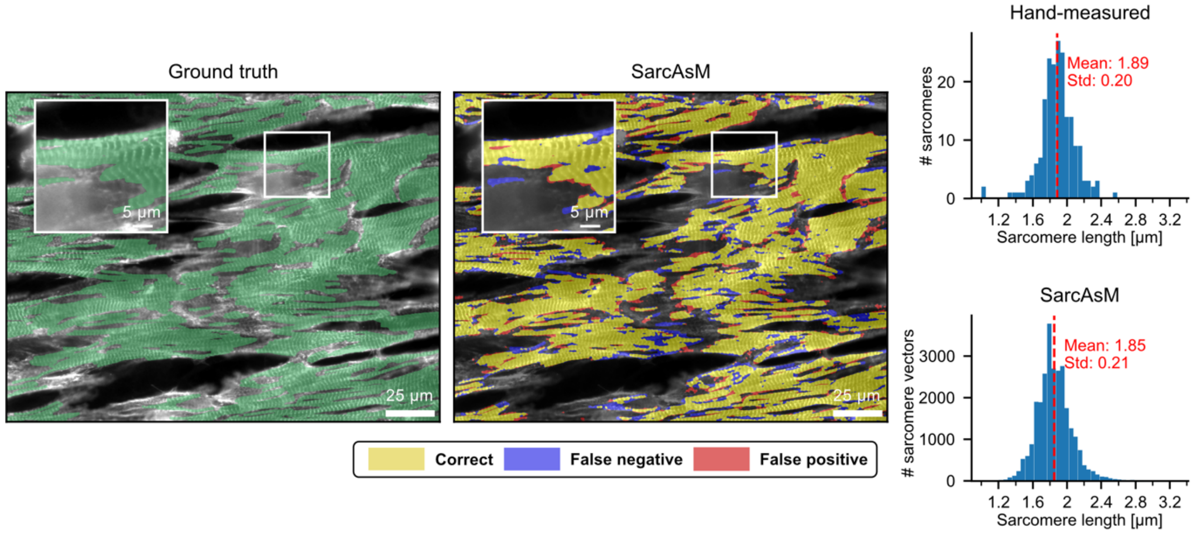


**Sample 2: Human myoblasts stained for sarcomeric α-actinin (from Morris et al. 2020[3]).** **Left panel**: Microscopy image with green overlay depicting the ground truth sarcomere mask. **Middle panel**: SarcAsM algorithm output displayed as a false color map of sarcomere predictions, where yellow indicates correct predictions, blue represents false negatives, and red shows false positives. Segmentation performance metrics: precision 0.90, recall 0.85, F1-score 0.87. **Right panel**: Comparative histograms of sarcomere length distributions obtained through manual hand-measurement and analysis with SarcAsM. Respective means and standard deviations are shown in red.

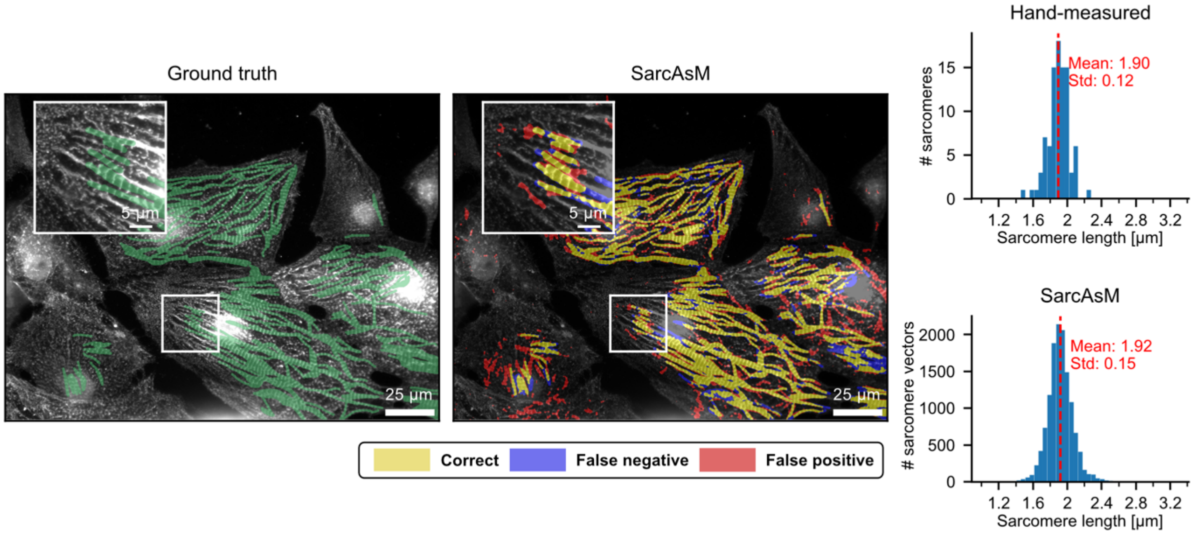


**Sample 3: hiPSC-derived cardiomyocyte stained for alpha-actinin 2 (own data). Left panel**: Microscopy image with green overlay depicting the ground truth sarcomere mask. **Middle panel**: SarcAsM algorithm output displayed as a false color map of sarcomere predictions, where yellow indicates correct predictions, blue represents false negatives, and red shows false positives. Segmentation performance metrics: precision 0.68, recall 0.81, F1-score 0.74. **Right panel**: Comparative histograms of sarcomere length distributions obtained through manual hand-measurement and analysis with SarcAsM. Respective means and standard deviations are shown in red.

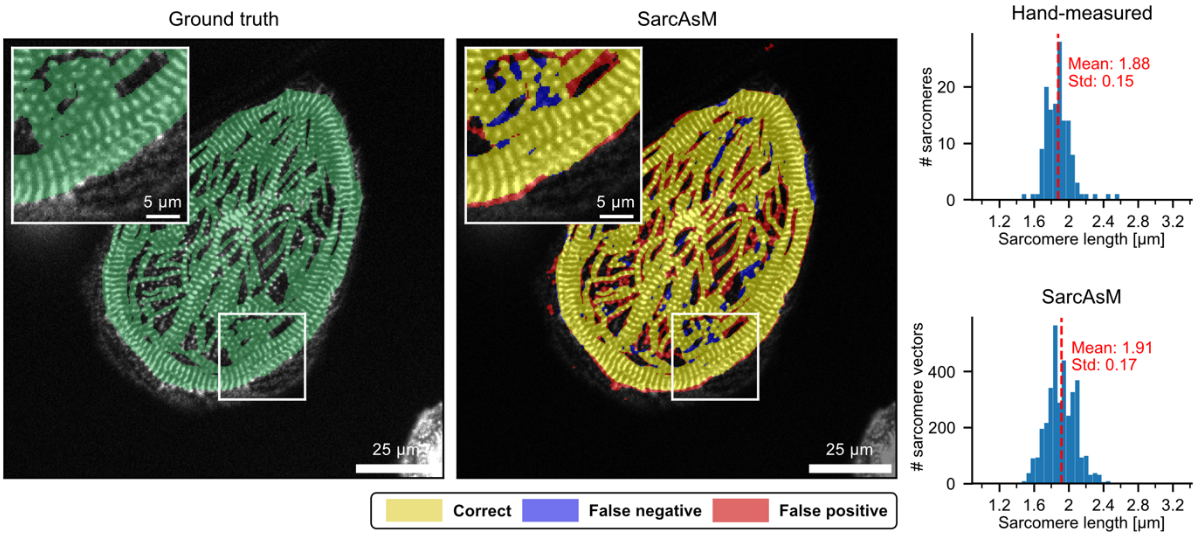


**Sample 4: hiPSC-derived CMs with mAppleACTN-2 lentivirus transduction (from Zhao et al. 2021)[4]. Left panel**: Microscopy image with green overlay depicting the ground truth sarcomere mask. **Middle panel**: SarcAsM algorithm output displayed as a false color map of sarcomere predictions, where yellow indicates correct predictions, blue represents false negatives, and red shows false positives. Segmentation performance metrics: precision 0.83, recall 0.95, F1-score 0.88. **Right panel**: Comparative histograms of sarcomere length distributions obtained through manual hand-measurement and analysis with SarcAsM. Respective means and standard deviations are shown in red.

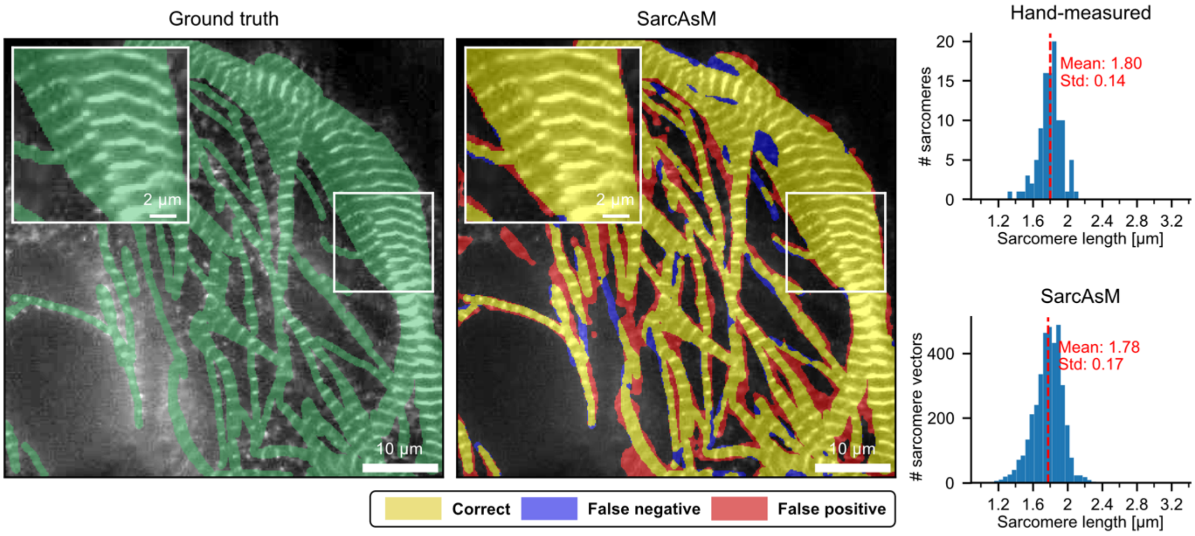


**Sample 5: hiPSC-derived TTN-GFP cardiomyocyte in 2D culture (from Zhao et al. 2021[4]).** **Left panel**: Microscopy image with green overlay depicting the ground truth sarcomere mask. **Middle panel**: SarcAsM algorithm output displayed as a false color map of sarcomere predictions, where yellow indicates correct predictions, blue represents false negatives, and red shows false positives. Segmentation performance metrics: precision 0.78, recall 0.92, F1-score 0.84. **Right panel**: Comparative histograms of sarcomere length distributions obtained through manual hand-measurement and analysis with SarcAsM. Respective means and standard deviations are shown in red.

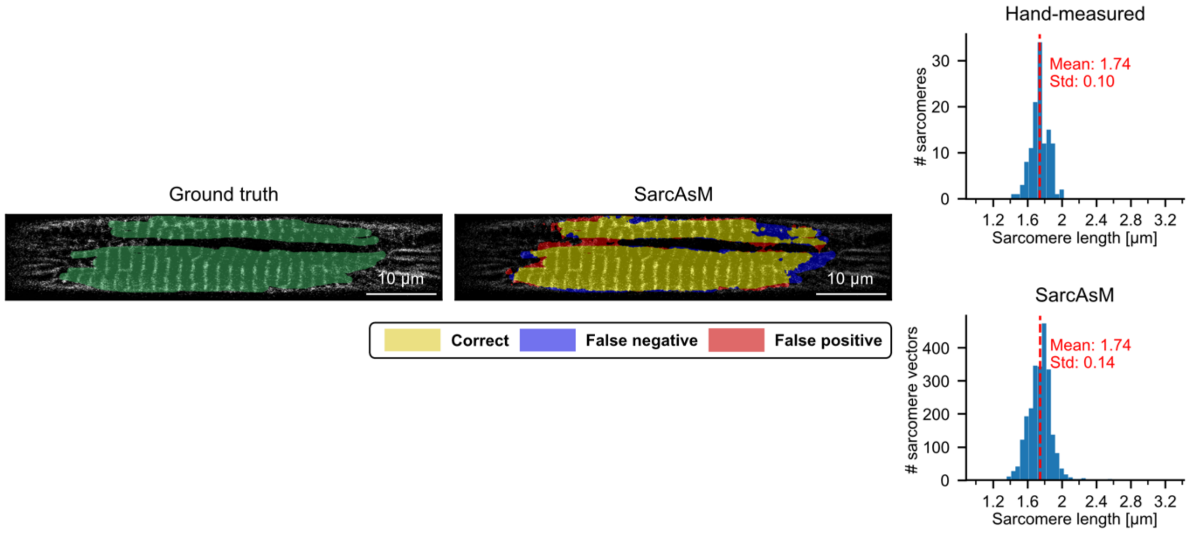


**Sample 6: Micropatterned hiPSC-derived ACTN2-citrine cardiomyocyte (own data). Left panel**: Microscopy image with green overlay depicting the ground truth sarcomere mask. **Middle panel**: SarcAsM algorithm output displayed as a false color map of sarcomere predictions, where yellow indicates correct predictions, blue represents false negatives, and red shows false positives. Segmentation performance metrics: precision 0.91, recall 0.88, F1-score 0.90. **Right panel**: Comparative histograms of sarcomere length distributions obtained through manual hand-measurement and analysis with SarcAsM. Respective means and standard deviations are shown in red.

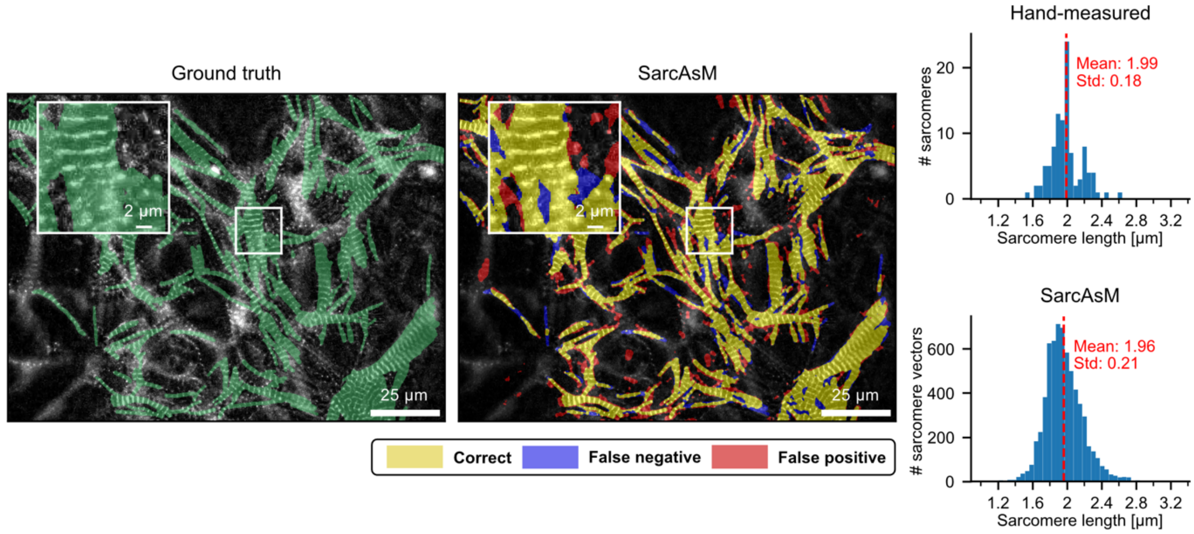


**Sample 7: hiPSC-derived ACTN2-GFP cardiomyocyte in 2D culture (from Sharma et al. 2018[5]). Left panel**: Microscopy image with green overlay depicting the ground truth sarcomere mask. **Middle panel**: SarcAsM algorithm output displayed as a false color map of sarcomere predictions, where yellow indicates correct predictions, blue represents false negatives, and red shows false positives. Segmentation performance metrics: precision 0.76, recall 0.85, F1-score 0.80. **Right panel**: Comparative histograms of sarcomere length distributions obtained through manual hand-measurement and analysis with SarcAsM. Respective means and standard deviations are shown in red.

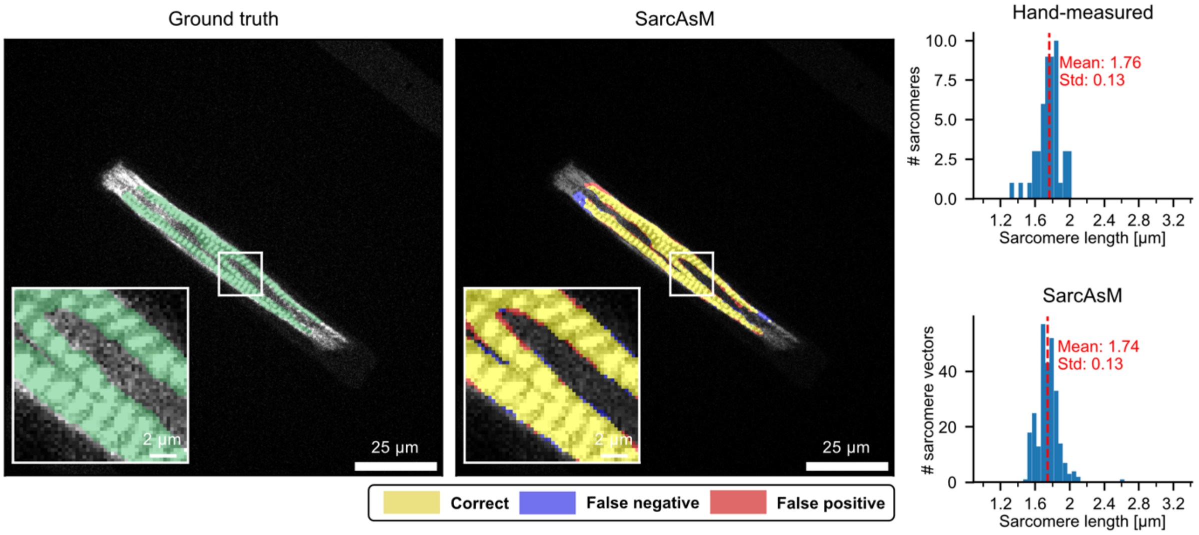


**Sample 8: Micropatterned hiPSC-derived cardiomyocyte (from Zhao et al. 2021[4]). Left panel**: Microscopy image with green overlay depicting the ground truth sarcomere mask. **Middle panel**: SarcAsM algorithm output displayed as a false color map of sarcomere predictions, where yellow indicates correct predictions, blue represents false negatives, and red shows false positives. Segmentation performance metrics: precision 0.89, recall 0.91, F1-score 0.90. **Right panel**: Comparative histograms of sarcomere length distributions obtained through manual hand-measurement and analysis with SarcAsM. Respective means and standard deviations are shown in red.

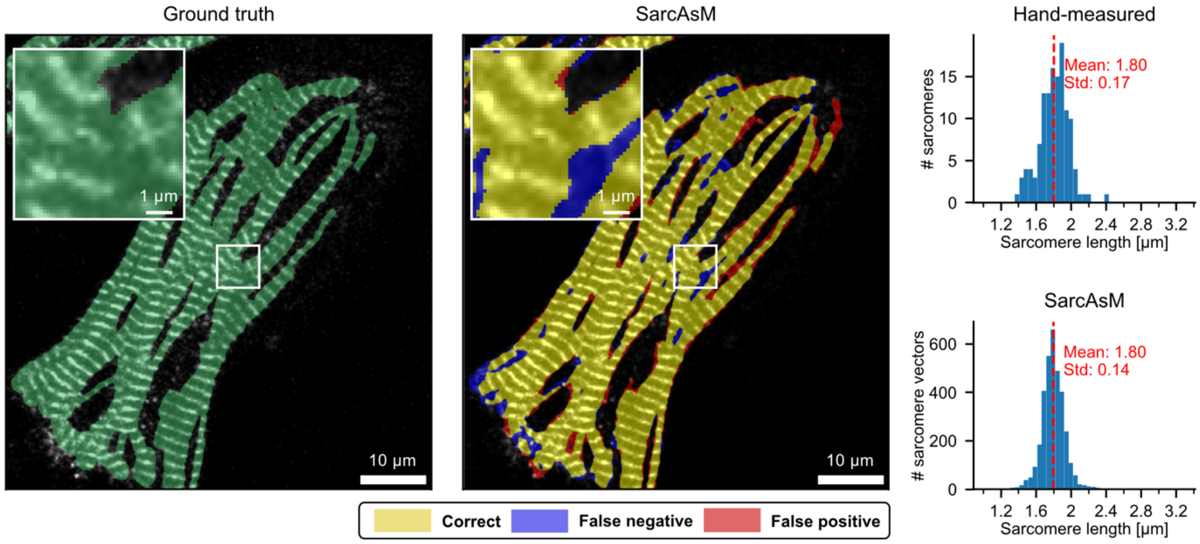


**Sample 9: TTN-GFP hiPSC-derived cardiomyocyte (own data). Left panel**: Microscopy image with green overlay depicting the ground truth sarcomere mask. **Middle panel**: SarcAsM algorithm output displayed as a false color map of sarcomere predictions, where yellow indicates correct predictions, blue represents false negatives, and red shows false positives. Segmentation performance metrics: precision 0.91, recall 0.91, F1-score 0.91. **Right panel**: Comparative histograms of sarcomere length distributions obtained through manual hand-measurement and analysis with SarcAsM. Respective means and standard deviations are shown in red.

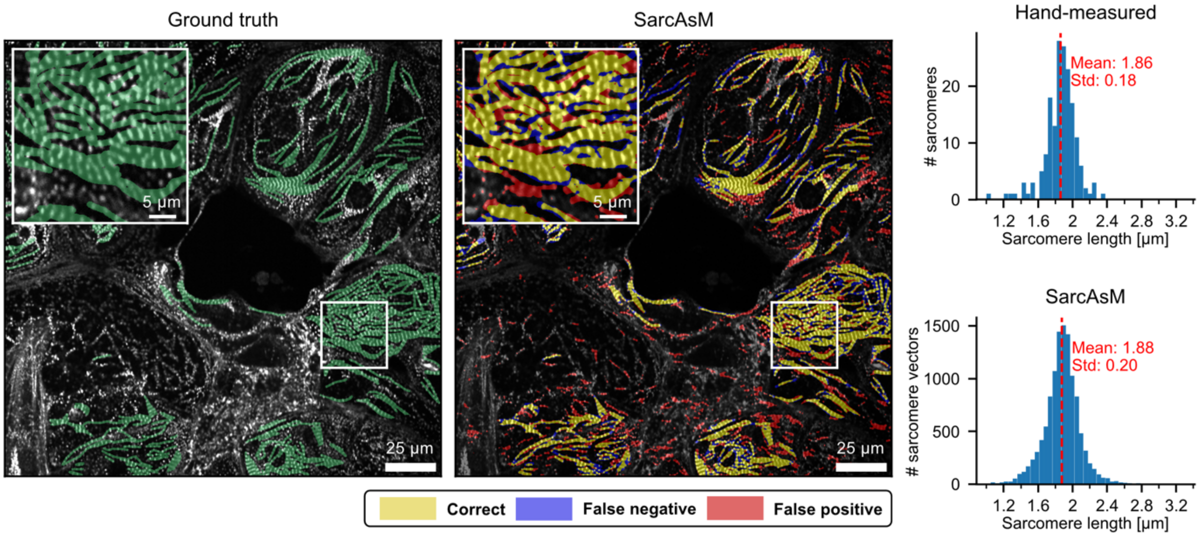


**Sample 10: 2D cultured ACTN2-citrine hiPSC-derived cardiomyocytes treated with 1 µM mavacamten (own data). Left panel**: Microscopy image with green overlay depicting the ground truth sarcomere mask. **Middle panel**: SarcAsM algorithm output displayed as a false color map of sarcomere predictions, where yellow indicates correct predictions, blue represents false negatives, and red shows false positives. Segmentation performance metrics: precision 0.63, recall 0.73, F1-score 0.68. **Right panel**: Comparative histograms of sarcomere length distributions obtained through manual hand-measurement and analysis with SarcAsM. Respective means and standard deviations are shown in red.

### SI - Supplementary Data

All training data, test data, benchmarking data, exemplary data of chronic and acute drug screening experiments, and pre-trained neural network models are available on Zenodo (DOI: 10.5281/zenodo.8232838).

SarcAsM is available on GitHub (https://github.com/danihae/SarcAsM) and Python Package Index (https://pypi.org/project/sarc-asm/). The published version of the code is also archived on Zenodo (DOI: 10.5281/zenodo.13925145), while the GitHub repository will always contain the most up-to-date version. SarcAsM has detailed online documentation with instructions for installation and API usage (https://sarcasm.readthedocs.io/en/).

For 3D U-Net, regular U-Net and U-Net++, our custom implementations are used (https://github.com/danihae/bio-image-unet).

**SI – Supplementary Figures**

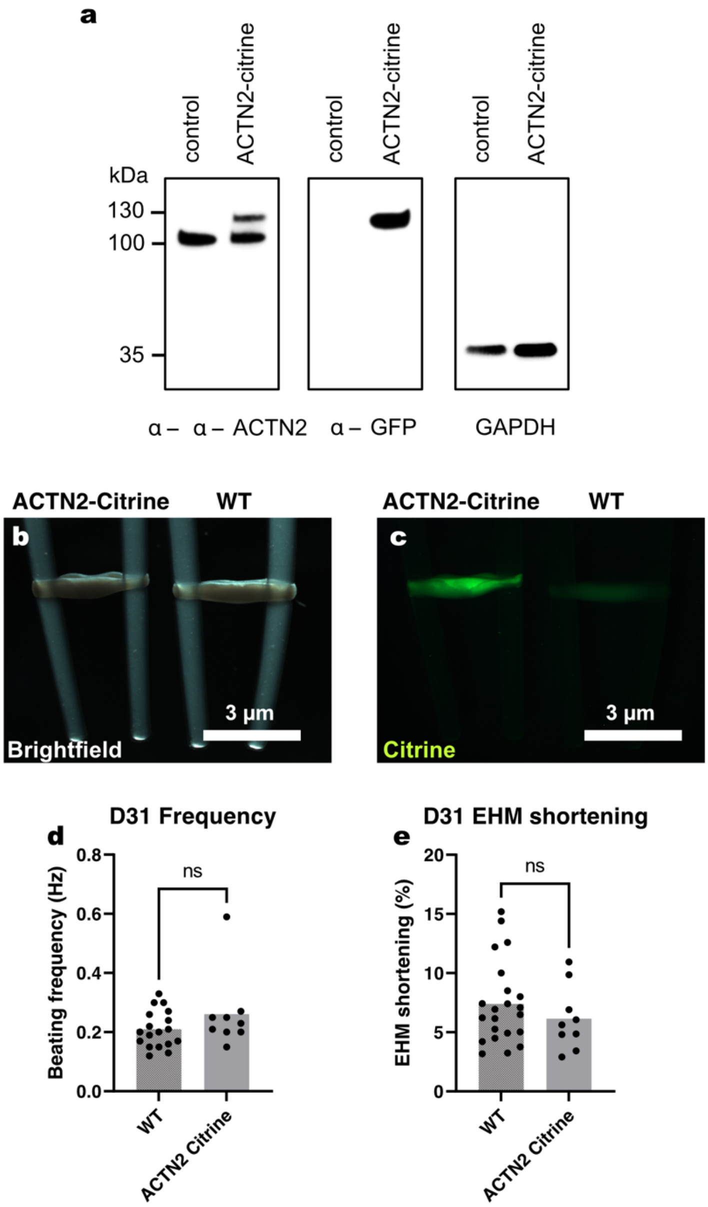


**Supplementary Figure 1: Characterization and validation of ACTN2-citrine Z-band reporter model**. (**a**) Western-blot analysis of ACTN2 and citrine expression in hiPSC-derived cardiomyocytes from control (not edited) and edited (ACTN2-citrine) lines. Total cell lysates were separated by SDS-PAGE, transferred onto a PVDF membrane, and stained with anti-ACTN2 and anti-GFP antibodies. Additional staining of the same PVDF membrane after washing with anti-GAPDH antibodies confirmed similar loading. Molecular mass markers in kDa are indicated on the left. Brightfield (**b**) and fluorescence (**c**) images of engineered human myocardium (EHM – culture day 31) constructed from ACTN2-citrine or wildtype (WT) cardiomyocytes showed no morphological differences and the anticipated strong fluorescence signal in ACTN2-citrine EHM. Comparison of spontaneous beating frequencies (**d**) and twitch shortening (**e**), a correlate for force of contraction, showed no differences in contractile properties between ACTN2-citrine and WT EHM (n=22/9 for WT/ACTN2-citrine). Statistical significance was tested with unpaired, two-tailed Student t-test.

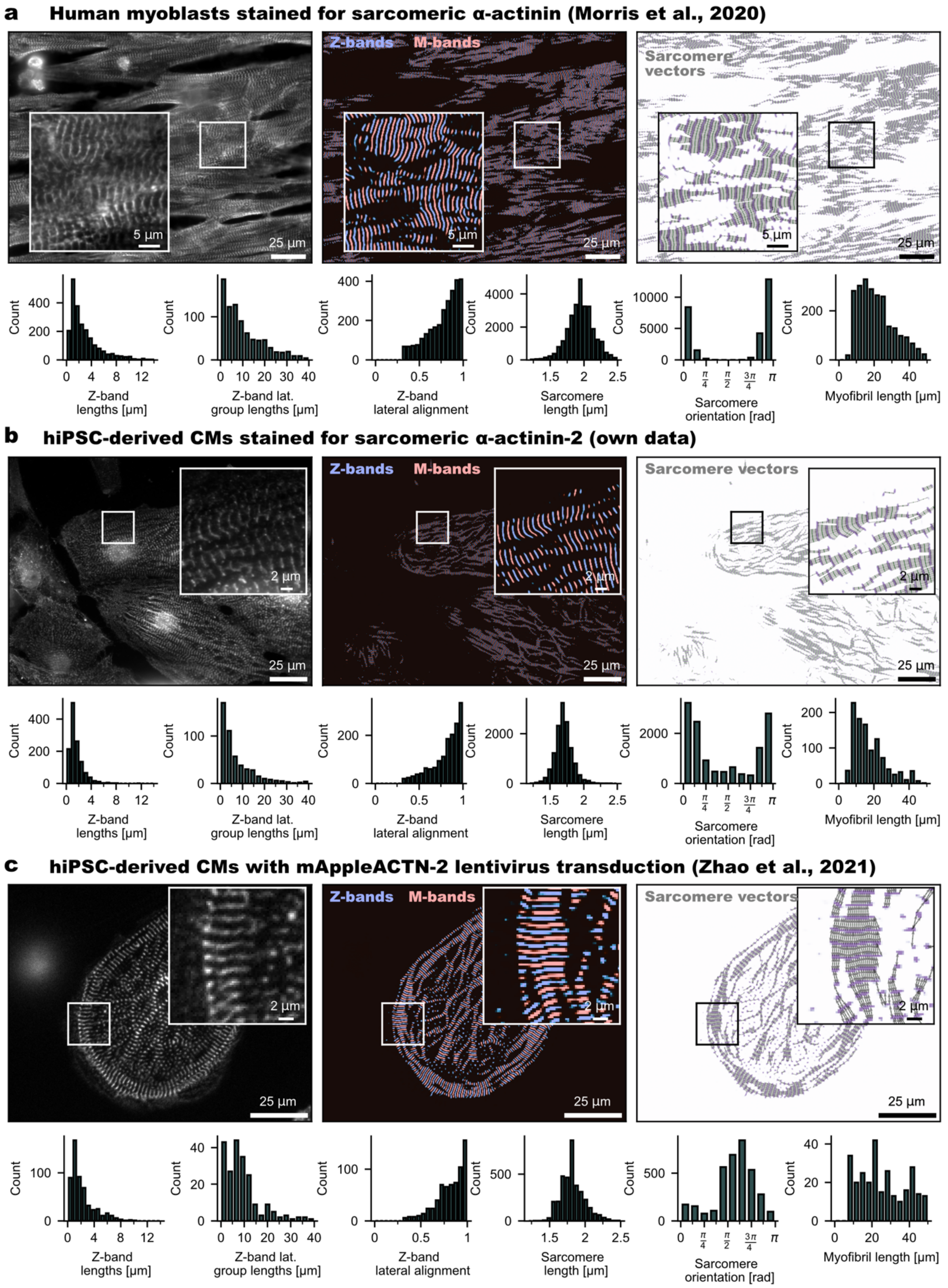


**Supplementary Figure 2: Application of SarcAsM to different data sets**. Fluorescence microscopy images and analysis results of (**a**) human skeletal muscle (https://doi.org/10.7280/D12Q2X)[3] and (**b**) hiPSC-CMs (own data) after immune-fluorescence labelling of ACTN2 (with additional labelling of nuclei [Hoechst] in **b**) as well as (**c**) hiPSC-CMs after lentiviral transduction of ACTN2-mApple (https://github.com/elejeune11/Sarc-Graph/blob/main/Supplementary_Information/E5.mp4). **Left**: input data. **Middle**: Z-bands and M-bands detected by deep learning. **Right**: sarcomere vectors (grey) and Z-bands (purple). **Bottoms**: Histograms of selected structural features.

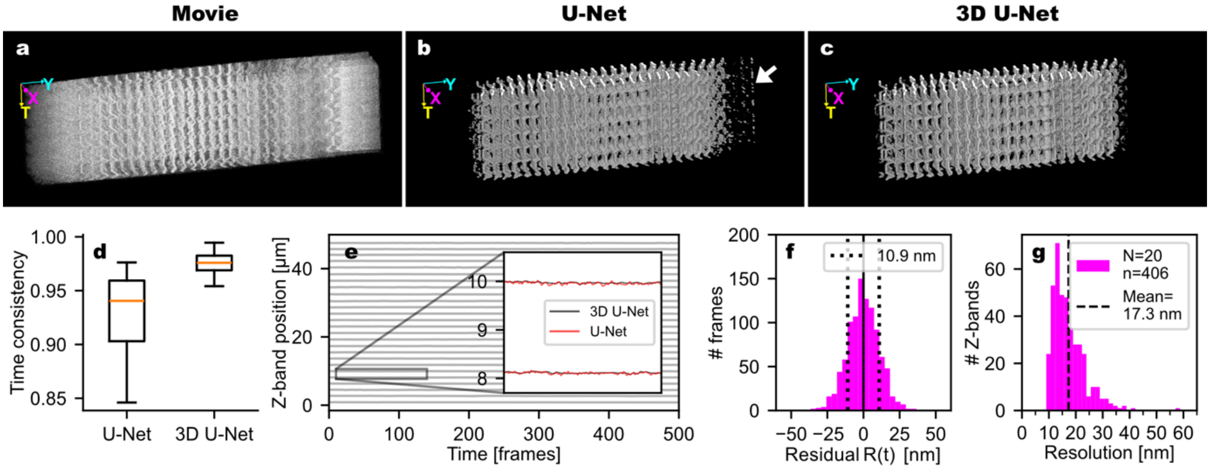


**Supplementary Figure 3: Validation of 3D U-Net for Z-band segmentation and tracking.** (**a**) 3D (T,X,Y) rendering of first 5 seconds of high-speed movie (66 frames per second) of ACTN2-citrine cardiomyocyte on micropatterned 20 kPa polyacrylamide gel. (**b**) 3D rendering of Z-bands segmented by standard U-Net of data in **a**, showing flickering and discontinuous Z-bands (white marker). (**c**) 3D rendering of Z-bands segmented by 3D U-Net of data in **a**, showing smooth and time-consistent segmentation with reduced flickering artifacts. Refer to **Supplementary Movie 1** for a direct comparison of **a-c**. Both U-Net and 3D U-Net were trained with the same training data. (**d**) Comparison of time consistency between standard U-Net and 3D U-Net, quantified by the ratio of the sum of areas of Z-band objects present in more than 64 consecutive frames to the total area of Z-bands in 20 randomly selected movies of 128 frames each. Boxes show quartiles, the red lines the median and whisker the 5$^{th}$ and 95$^{th}$ percentile. (**e**) Z-band trajectories of LOI of non-contracting cardiomyocyte tracked based on standard U-Net and 3D U-Net, showing more less noisy trajectories for 3D U-Net (see inset). (**f**) Histogram of residuals *R* for quantification of fluctuation of each Z-band around its respective mean length. The standard deviation of *R* is a measure for the accuracy of tracking. (**g**) Tracking accuracies of 406 individual Z-band trajectories recorded from 20 representative quiescent cardiomyocytes.

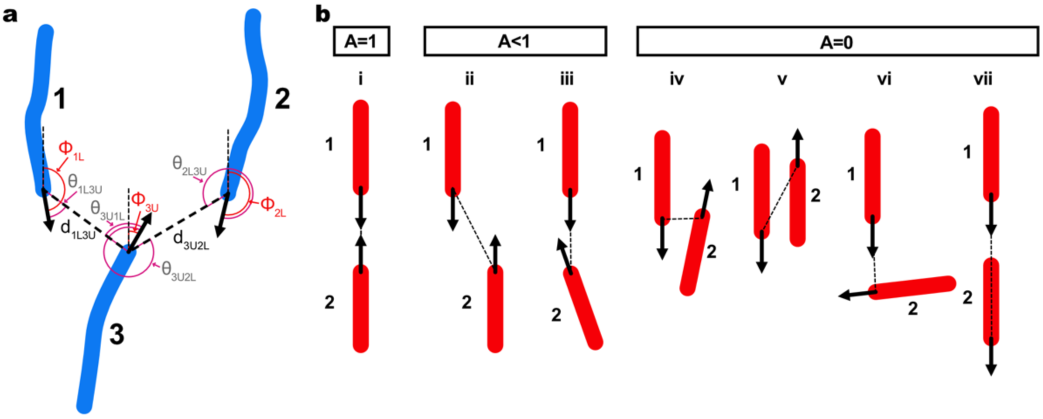


**Supplementary Figure 4: Z-band lateral connections and sarcomere domain analysis.** (**a**) Illustration of distances (d; thick dashed lines), orientations (Φ) and orientation differences (θ) of nearest neighbor Z-band (indices 1 to 3) ends (U=upper end, L=lower end). (**b**) Illustration of lateral alignment metric A of Z-band pairs (1 and 2 in red): (i) $A = 1$ for perfect antiparallel alignment ($\theta_{1L2U} = \theta_{2U1L} = 0$. $|\Phi 1L - \Phi 2U| = \pi$); $A < 1$ when the two Z-bands (ii) have an offset ($\theta_{1L2U} \neq 0$ and/or $\theta_{2U1L} \neq 0$) and/or (iii) or are additionally not oppositely oriented ($|\Phi 1L - \Phi 2U| \neq \pi$); A=0 (no alignment) when the two Z-ends are too close or apart (d<0.25 µm or d>5 µm), (iv,v) are interdigitating ($|\theta_{1L2U}| \geq \pi$ *and/or* $|\theta_{2U1L}| \geq \pi$), (vi, vii) or similarly oriented ($|\Phi 1L - \Phi 2U + \pi| \geq \pi/2$). 2π (rad) = 360°.

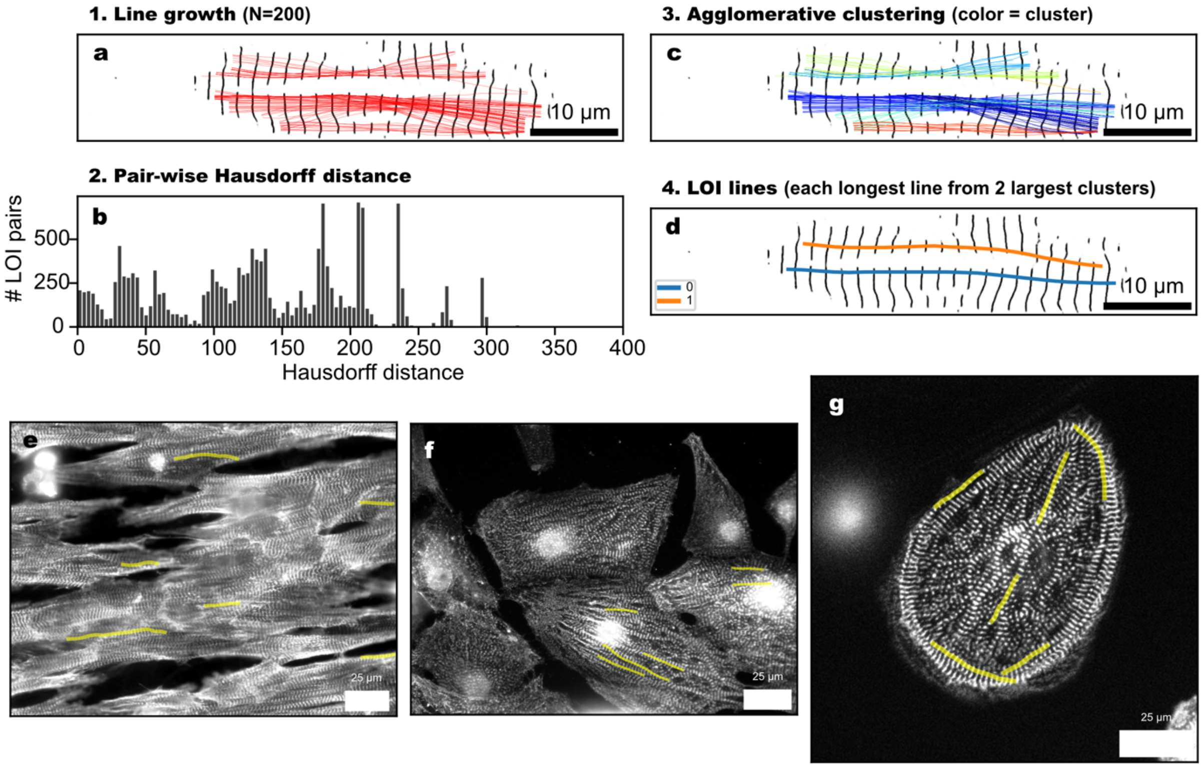


**Supplementary Figure 5: Automated line-of-interest (LOI) detection for sarcomere motion tracking.** (**a**) Red lines show result of line growth algorithm for persistence ($P = 8$) and minimal length $N_{min} = 10$. In the background the U-Net result is shown. (**b**) Pairwise Hausdorff distance between lines in **a**. (**c**) Result of agglomerative clustering of lines with pre-computed Hausdorff distances and threshold distance $d_{max} = 40$. Clustering lines are shown in same color. (**d**) Longest lines from two largest clusters are selected as LOI lines. Alternatively, one random line per cluster can be selected, or a straight line can be fitted to all points of a cluster. In the presented cell, the longest LOIs from the two largest clusters are shown. (**e-f**) LOI detection in three different samples using default parameters ($P = 8$, $d_{\max} = 40$), showing each one random line from the 6 largest clusters. Examples for automated LOI detection in the following data sources: **e** human skeletal muscle cells stained for alpha-actinin 2 (https://doi.org/10.7280/D12Q2X); **f** hiPSC-derived cardiomyocytes stained for alpha-actinin 2 (own data); **g** hiPSC-derived cardiomyocyte with mAppleACTN-2 lentivirus transduction (https://github.com/Sarc-Graph/sarcgraph).

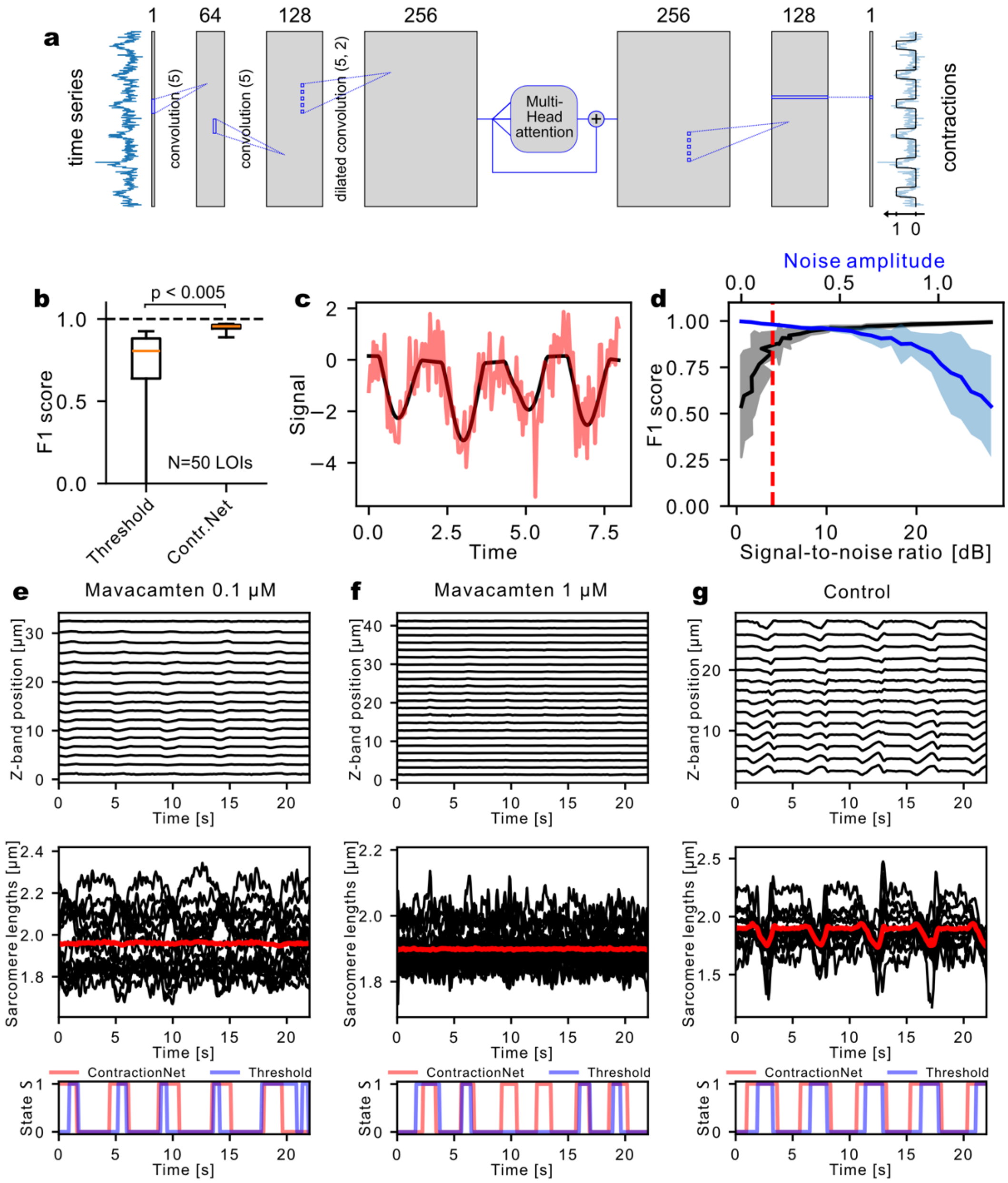


**Supplementary Figure 6: Network architecture and validation of ContractionNet.** (**a**) Schematic of ContractionNet. (**b**) Validation of ContractionNet using real data of a representative set of 50 LOIs, compared against threshold-based method (deviation of mean sarcomere length from baseline (maximum of histogram of lengths) by more than threshold of -0.01 µm). Boxes show quartiles, the red line the median and whiskers the 5$^{th}$ and 95$^{th}$ percentile. (**c**) Simulated time-series mimicking cardiac contraction data, with differing amplitudes and baseline shift (black). Red line shows data with additional normal-distributed noise with amplitude 0.15. (**d**) Validation of ContractionNet using simulated data, as seen in **c**. For time-series with different noise-amplitudes/signal-to-noise ratios (SNRs) the F1-score was calculated between the ContractionNet output and the known ground truth. At high noise amplitudes, corresponding to small SNRs (e.g. SNR of 3 corresponding to data in **c**, see red dashed line), ContractionNet was still able to infer the contraction state with high accuracy (F1-score>0.9). Examples of contraction interval analyses using ContractionNet and threshold-based methods for 3 representative LOIs (0.1 µM [**e**] and 1 µM [**f**] Mavacamten both with weak contractions and control [**g**] with strong regular contractions; Z-band segmentation with regular 2D U-Net). Red lines in second row show average sarcomere length, thin black lines show individual sarcomere lengths.

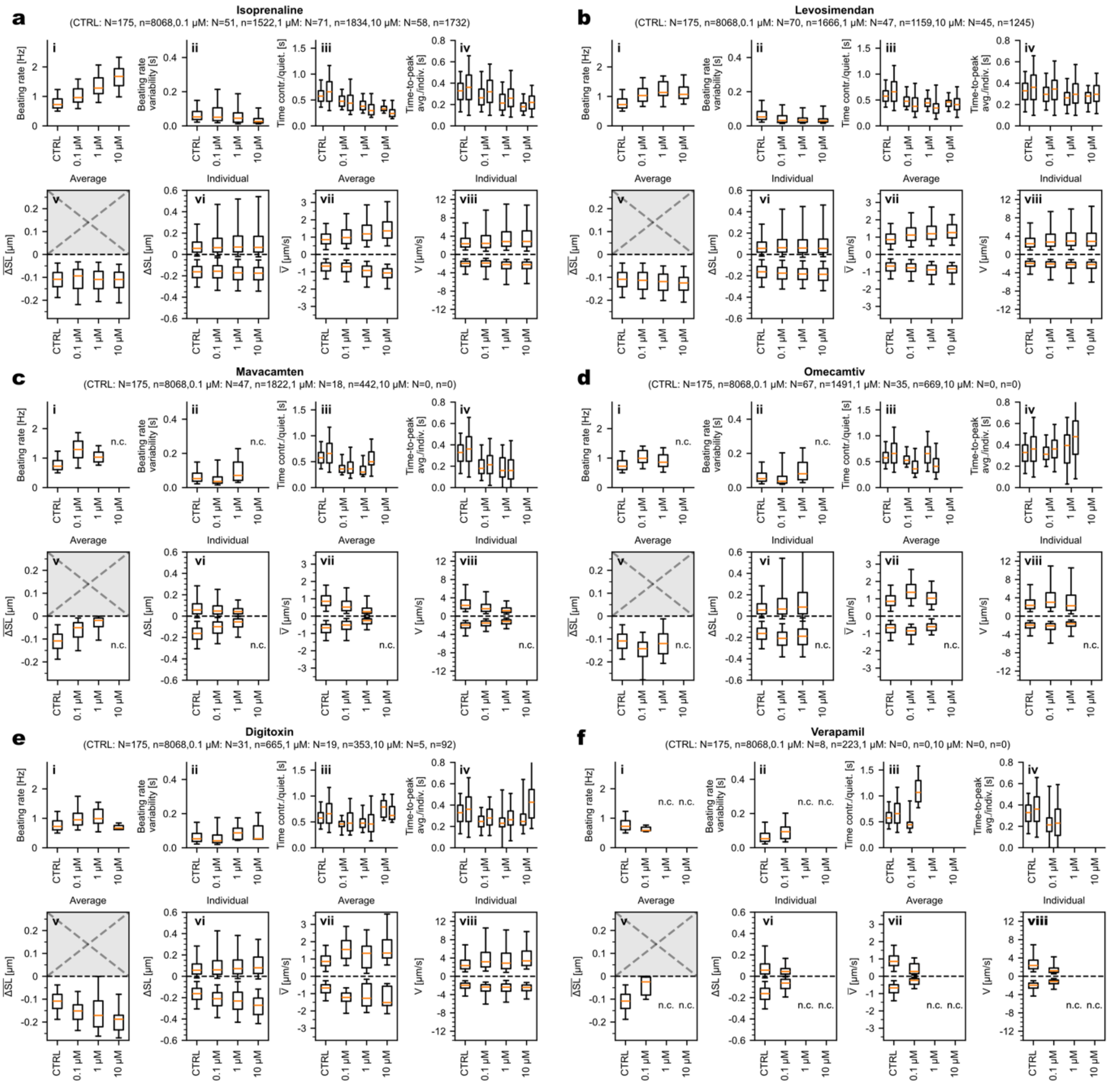


**Supplementary Figure 7: Acute drug effects on individual and average sarcomere dynamics**. Summary of obtained data after exposure of cardiomyocytes to (**a**) Isoprenaline, (**b**) Levosimendan, (**c**) Mavacamten, (**d**) Omecamtiv, (**e**) Digitoxin, and (**f**) Verapamil (**data related to main Figure 6**). The following parameters were documented for each compound: (**i**) beating rate at different drug concentrations; (**ii**) beating rate variability, quantified as standard deviation of time differences between start time point of consecutive contraction intervals; (**iii**) duration of contraction intervals (left boxplot) and quiescent intervals between contractions (right boxplot); (**iv**) time-to-peak (maximum shortening) of average sarcomeres (left) and individual sarcomeres (right); (**v**) maximum shortening of sarcomere average per LOI; (**vi**) maximal shortening (lower) and lengthening (upper) amplitudes of individual sarcomeres; (**vii**) maximum shortening (lower) and lengthening (upper) velocities of sarcomere average per LOI; (**viii**) maximal shortening (lower) and elongation (upper) velocities of individual sarcomeres. Boxes show the quartiles, the red line the median, and whiskers the 5$^{th}$ and 95$^{th}$ percentile. N=number of LOIs (max. 2 per cell); n=number of evaluated individual contractions (number sarcomeres x number contractions).

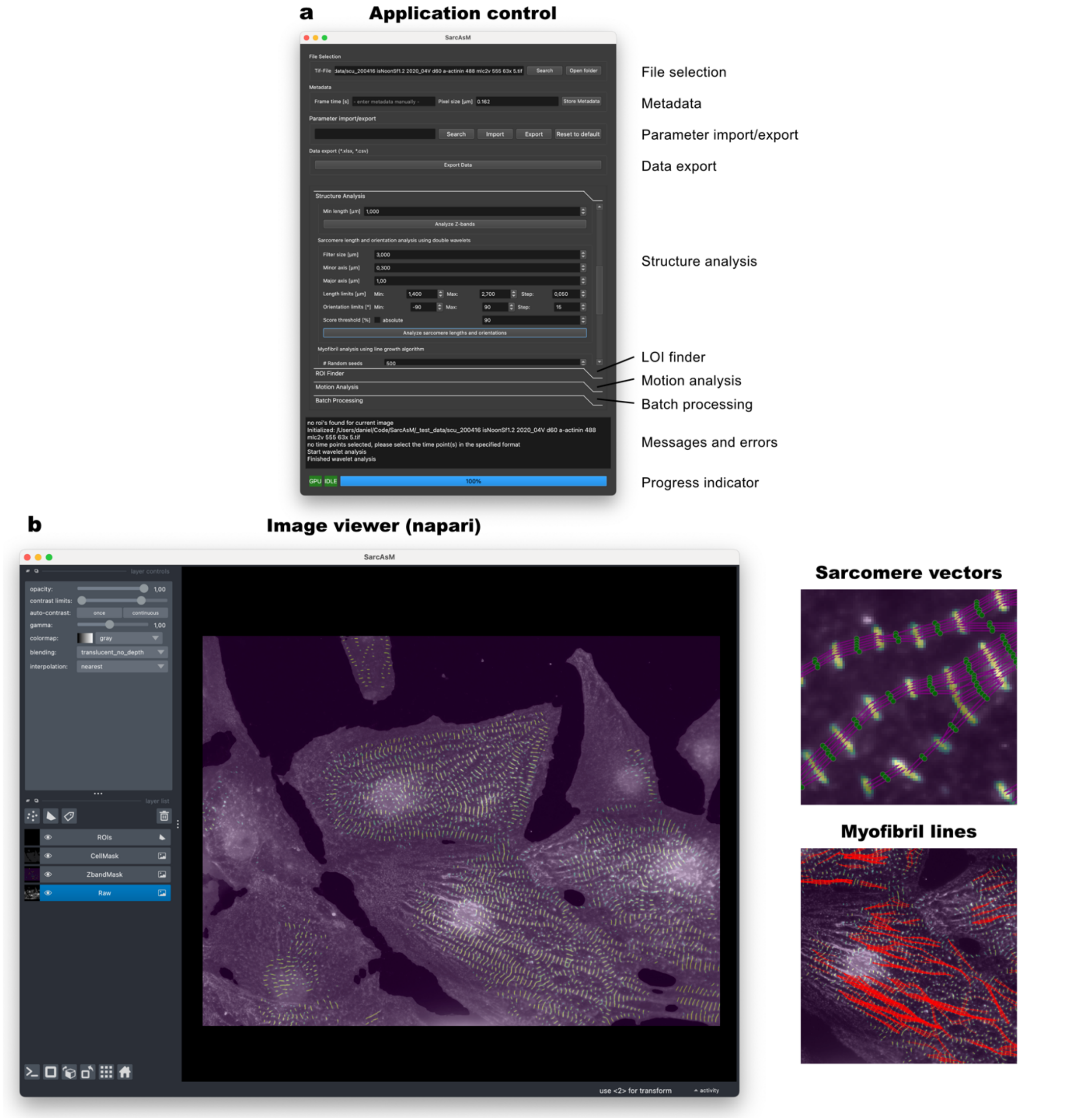


**Supplementary Figure 8: SarcAsM graphical user interface of standalone application.** (**a**) Application control user interface with file and metadata handling, parameter export/import, data export, all functions of SarcAsM (structure analysis, LOI finder, motion analysis), batch processing mode, message field, and progress indicator. (**b**) Image viewer, based on Napari. Left: Image with yellow overlay showing sarcomere Z-bands. Right: sarcomere vectors and myofibrils as displayed in image viewer upon completed analysis.

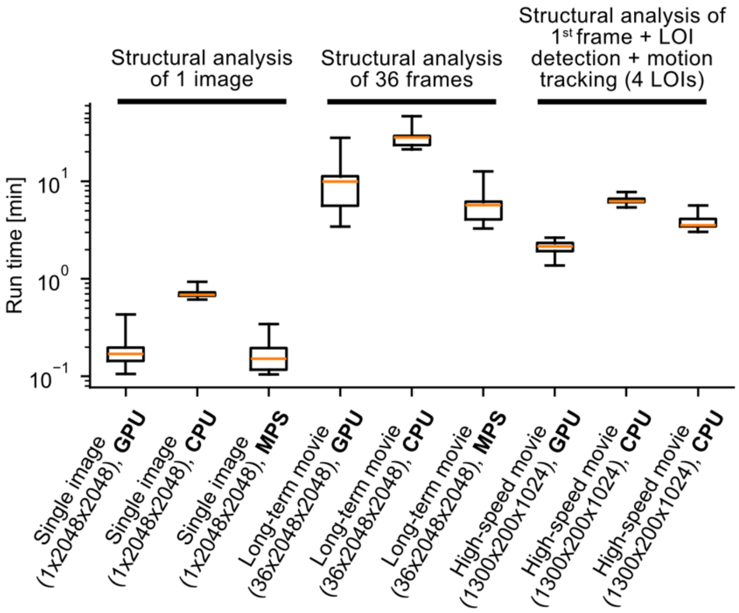


**Supplementary Figure 9: SarcAsM run times for three analytical scenarios across different computational platforms.** Tests were performed on a workstation GPU (NVIDIA Quadro RTX 5000, Intel Xeon W-2245), a standard notebook CPU (Intel Core i7-10750H), and a MacBook Pro (2022 with M1 Pro using Metal Performance Shaders (MPS) with GPU-like capabilities). Scenarios include structural analysis of single images (2048×2048 pixels), long-term movies with 36 frames (36×2048×2048 pixels), and high-speed recordings with first frame analysis, LOI detection and motion tracking for 4 LOIs (1500×2048×2048 pixels). Box plots show quartiles with median (red line), and whiskers indicate minimum/maximum values. The y-axis uses a logarithmic scale for run times in minutes.

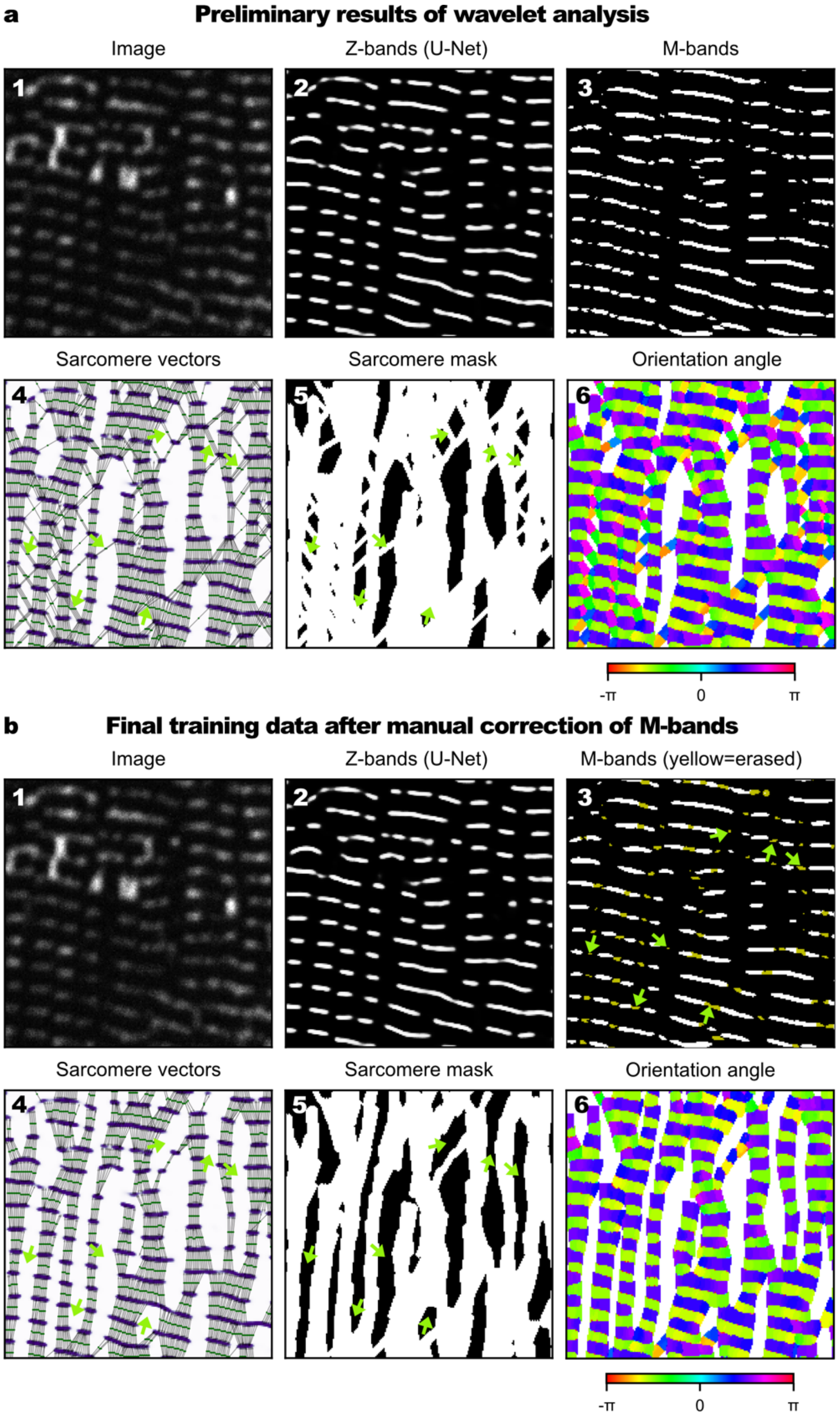


**Supplementary Figure 10: Training data generation for sarcomere M-bands, sarcomere orientation and sarcomere masks**. (**a**) Preliminary output of double wavelet analysis (details in **Supplementary Note 1**). (1) Representative section of training image (confocal image of ACTN2-citrine cardiomyocyte). (2) Z-band mask predicted by U-Net trained on manually annotated Z-bands. (3) M-band mask output by wavelet analysis. Preliminary (4) sarcomere vectors with multiple false positives (yellow arrows highlight examples), (5) sarcomere mask, and (6) sarcomere orientation angle map. (**b**) Final training data after manual correction of the M-band mask. (1) and (2) as above. (3) Final M-band mask with erased regions shown in yellow. Final (4) sarcomere vectors after correction of M-bands, (5) sarcomere mask, and (6) sarcomere orientation angle map. Yellow arrows highlight previously false-positive sarcomeres that were corrected.

### SI – Supplementary Tables

| | |
|---|---|
| ACTN2 wild type | 5’ – TGGATTACGCTGCGTTCTCTTCCGCACTCTACGGGGAGAGCGATCTG**TGA** – 3’ |
| ACTN2 edited | 5’ – TGGATTACGCTGCGTTCTCTTCTGCTTTGTATGGGGAGAGCGATCTG*GGAGGTGGAGGTGGAGCT*GTGAGCAAGGGCGAGGAGCTGTTCACCGGGGTGGTGCCCATCCTGGTCGAGCTGGACGGCGACGTAAACGGCCACAAGTTCAGCGTGTCCGGCGAGGGCGAGGGCGATGCCACCTACGGCAAGCTGACCCTGAAGTTCATCTGCACCACCGGCAAGCTGCCCGTGCCCTGGCCCACCCTCGTGACCACCTTCGGCTACGGCCTGATGTGCTTCGCCCGCTACCCCGACCACATGAAGCAGCACGACTTCTTCAAGTCCGCCATGCCCGAAGGCTACGTCCAGGAGCGCACCATCTTCTTCAAGGACGACGGCAACTACAAGACCCGCGCCGAGGTGAAGTTCGAGGGCGACACCCTGGTGAACCGCATCGAGCTGAAGGGCATCGACTTCAAGGAGGACGGCAACATCCTGGGGCACAAGCTGGAGTACAACTACAACAGCCACAACGTCTATATCATGGCCGACAAGCAGAAGAACGGCATCAAGGCCAACTTCAAGATCCGCCACAACATCGAGGACGGCGGCGTGCAGCTCGCCGACCACTACCAGCAGAACACCCCCATCGGCGACGGCCCCGTGCTGCTGCCCGACAACCACTACCTGAGCTACCAGTCCAAGCTGAGCAAAGACCCCAACGAGAAGCGCGATCACATGGTCCTGCTGGAGTTCGTGACCGCCGCCGGGATCACTCTCGGCATGGACGAGCTATACAAG**TGA** – 3’ |

**Supplementary Table 1: Schematic representation of ACTN2 edited alleles.** Blue indicates gRNA target sequence, red is the PAM motif, green indicates silent SNPs, stop codon is shown in bold, linker sequence in italics. Monomeric citrine coding sequence (without start and stop codons in green) is abbreviated as citrine throughout manuscript (https://www.fpbase.org/protein/mcitrine/ with V163A & S175G substitutions for improved folding and brightness; Heim et al. 2007; PMID: 17259991).

| Dataset | Cell Type and Labeling | Acquisition Details | Source / Reference |
|---|---|---|---|
| ACTN2-Citrine-2D | hiPSC-derived cardiomyocytes in 2D culture, endogenous ACTN2-citrine tag | Yokogawa CV8000 spinning disk confocal microscope, 60x water immersion objective, 0.104 µm pixel size | Own data, details see methods section |
| ACTN2-Citrine-MPat | Individual hiPSC-derived cardiomyocytes on micropatterned polyacrylamide gel, endogeneous ACTN2-citrine tag, | Leica SP5 confocal microscope, 100x oil objective, 0.067 µm pixel size, single frames from 66 fps movies | Own data, details see methods section |
| SkeletalMuscle-ACTN2 | hiPSC-derived skeletal myocytes, α-Actinin immunostaining or endogenous ACTN2-citrine tag | 3i spinning disk confocal microscope, 40x water immersion objective, 0.163 µm pixel size | Own data; see Hofemeier et al., 2021[6], 2022[7], and Kim et al., 2024[8]4/29/25 8:44:00 PM |
| ACTN2-Immuno | hiPSC-derived cardiomyocytes in 2D culture, ACTN2 immunostaining | Axio Imager 2 Zeiss, 63x oil immersion objective, 0.114 µm pixel size | Own data, see Busley et al., 2024[9] |
| TTN-EGFP | hiPSC-derived cardiomyocytes in 2D culture, endogenous TTN-eGFP tag | Yokogawa CQ1 spinning disk or Molecular Devices ImageXpress Micro Confocal, both 60x air objective, 0.114 µm pixel size | Own data |
| MultiCell-ACTN2-Immuno | Mixed cell types, α-Actinin immunostaining | Olympus IX-83 microscope, 40x oil immersion, 0.167 µm pixel size | Morris et al., 2020[3] |
| ACTN2-mEGFP | hiPSC-derived cardiomyocytes with ACTN2-mEGFP tag | 3i spinning disk confocal microscope, 63x water immersion objective, 0.124 µm pixel size | Gerbin et al., 2021[10] |

**Supplementary Table 2: Cell type, labeling, and acquisition details of images used as training data for generalist sarcomere detection model.** This table summarizes the datasets, including cell types studied, labeling methods applied, imaging acquisition parameters, and references to their sources.

**SI – Supplementary Movies (mp4 movie files uploaded separately):**

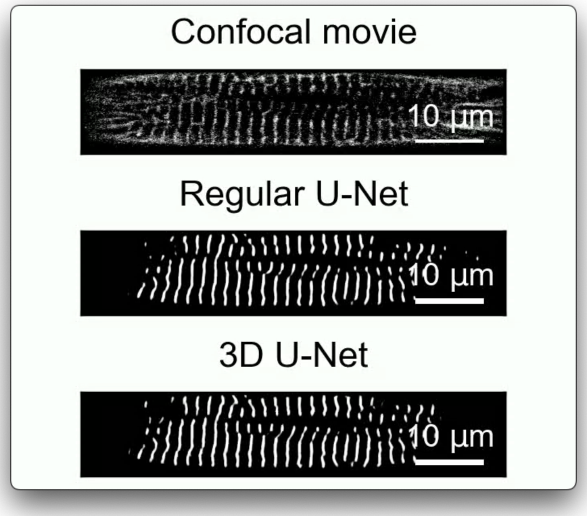


**Supplementary Movie 1: Detection of Z-bands from confocal microscopy movie (top) using U-Net neural network (middle) in comparison with time-consistent 3D-U-Net (bottom).** Panels show a representative single spontaneously beating ACTN2-citrine CM in real-time (66 frames/second) on micropatterned soft substrate (20 kPa Young's modulus). Both neural networks (regular U-Net and 3D U-Net) were trained on the same training data.

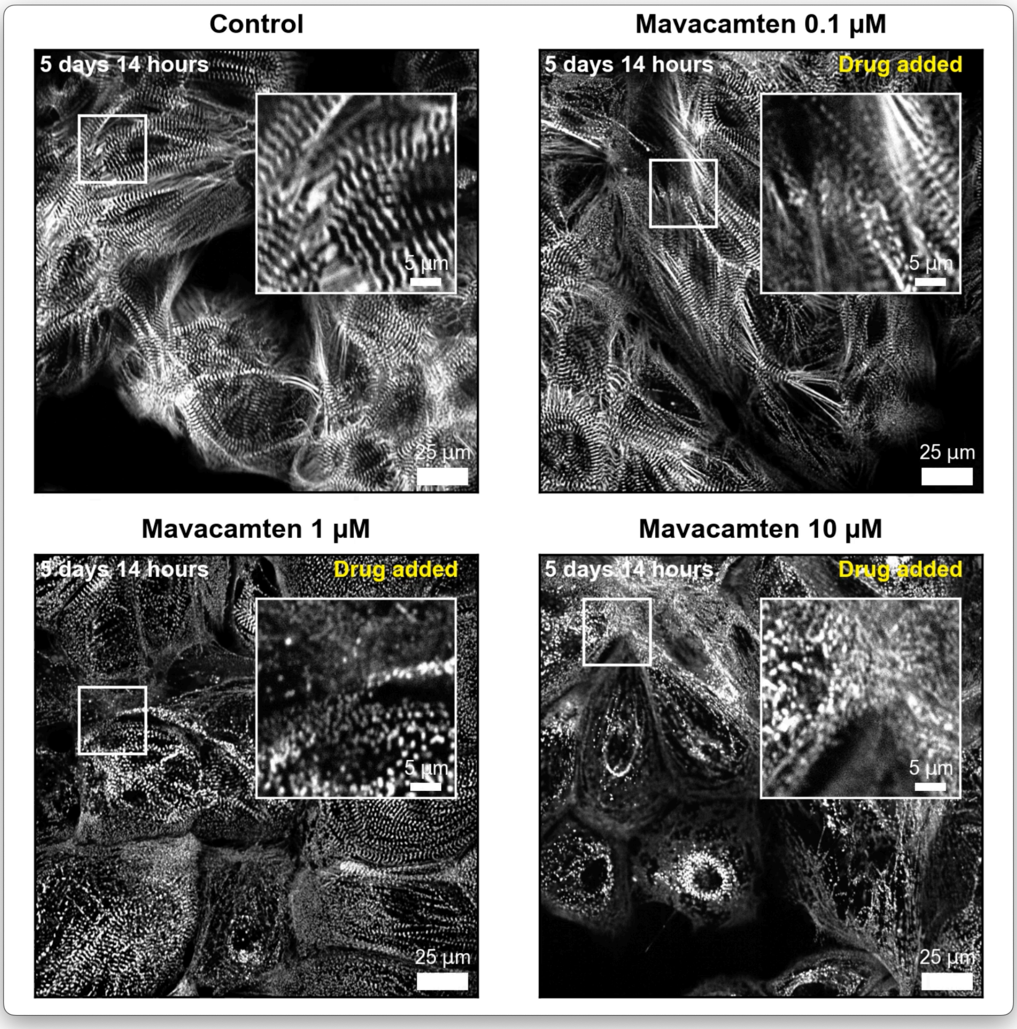


**Supplementary Movie 2: Time-course of chronic drug treatment of ACTN2-citrine CMs in unconstrained 2D culture in a 96-well plate.** Image acquisition was started after seeding of cells and continued for 8 days. For each condition, one representative field of view is shown.

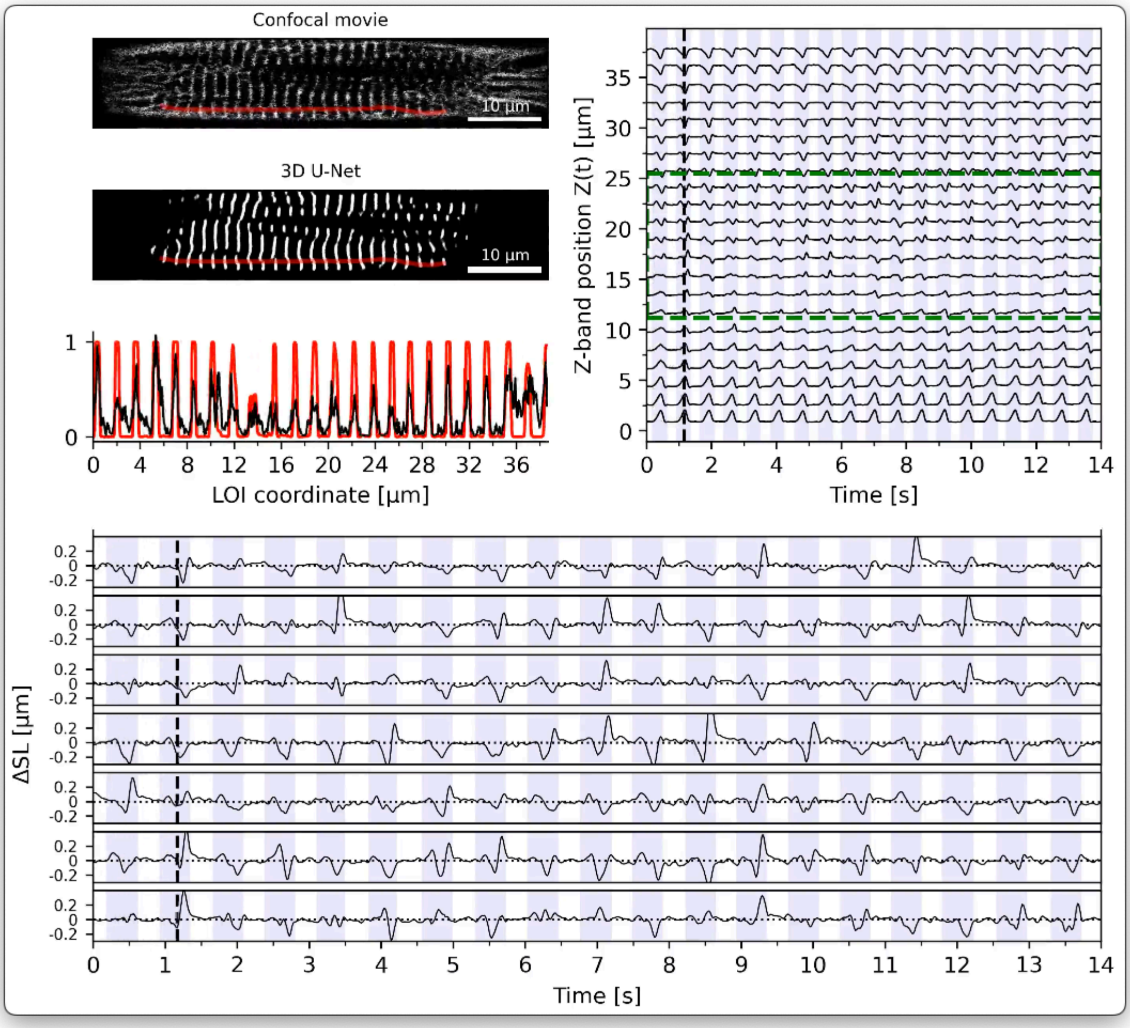


**Supplementary Movie 3: Animation of sarcomere motion tracking workflow.** Top left: real-time (66 frames/second) movie of representative spontaneously beating single ACTN2-citrine CM on micropatterned soft substrate (20 kPa Young's modulus) with sarcomere Z-bands predicted by 3D U-Net. Red line shows a line of interest (LOI) along which the motion of sarcomeres is tracked. Line profile along LOI for raw confocal data (black) and Z-bands (red). Top right: trajectories of individual sarcomere Z-bands localized from line profiles and tracked over time. Bottom: sarcomere length change of individual sarcomeres at highlighted LOI (top left).

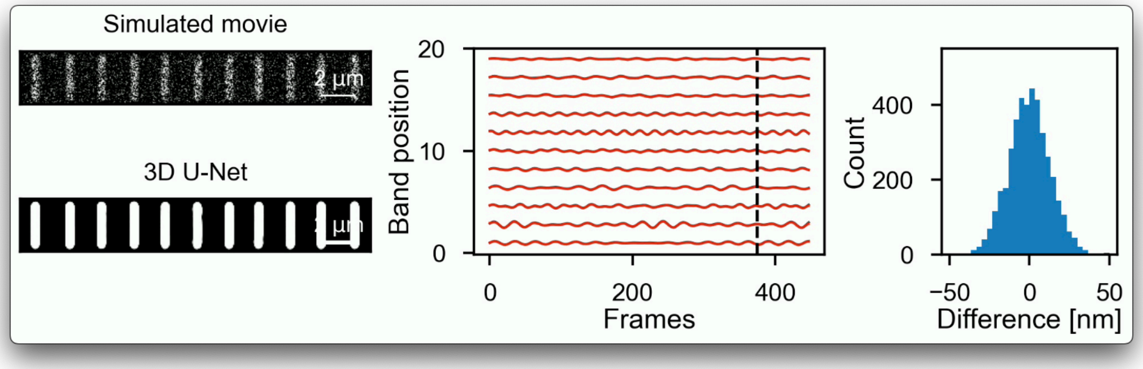


**Supplementary Movie 4: Analysis of simulated microscopy movie of rapidly moving sarcomere Z-bands.** Left: simulated high-noise microscopy movie with individual Z-bands following oscillatory motion with random frequencies and amplitudes and Z-bands reconstructed/predicted by 3D U-Net. Middle: tracked motion of individual Z-bands (red) and known input trajectories (black). Right: histogram of residual between tracked and input trajectories, with a standard deviation of 0.25 pixels, equivalent to 14 nm.
